# Biotic-response networks are an important organizer of the transcriptome in wild *Arabidopsis thaliana* populations

**DOI:** 10.64898/2026.03.11.711176

**Authors:** Ana Paula Leite Montalvão, Kevin Murray, Ilja Bezrukov, Natalie Betz, Lucas Henry, Paloma Duran, Patrick Boppert, Martina Kolb, TEAM PATHOCOM, Fabrice Roux, Joy Bergelson, Wei Yuan, Detlef Weigel

**Author notes:** correspondence (W.Y.), (D.W.). equal contribution (Ana Paula Leite Montalvão, Kevin Murray, Ilja Bezrukov).

## Abstract

Extensive laboratory experimentation has revealed conserved molecular pathways controlling growth and stress responses in plants, yet how these programs operate in natural settings remains poorly understood. We investigated transcriptome organization in wild populations of *Arabidopsis thaliana* by sampling plants from 60 natural sites in Europe and North America across two seasons. Transcriptomes varied extensively among individuals and showed largely continuous rather than discrete structure across geography and season. Although disease and microbial colonization were common in the wild, wild transcriptomes did not simply recapitulate canonical laboratory stress signatures. Measured microbial infection, environmental, and phenotypic variables explained only a modest fraction of total expression variation, but infection-associated signals accounted for the largest share of the explainable component. Consistent with this, biotic-response networks defined in controlled laboratory experiments were well conserved in wild transcriptomes, whereas control and abiotic-response networks were substantially reorganized. Together, these results suggest that while core transcriptional modules remain recognizable across environments, regulatory relationships among modules differ markedly between laboratory and natural contexts.

**Graphical Abstract:** 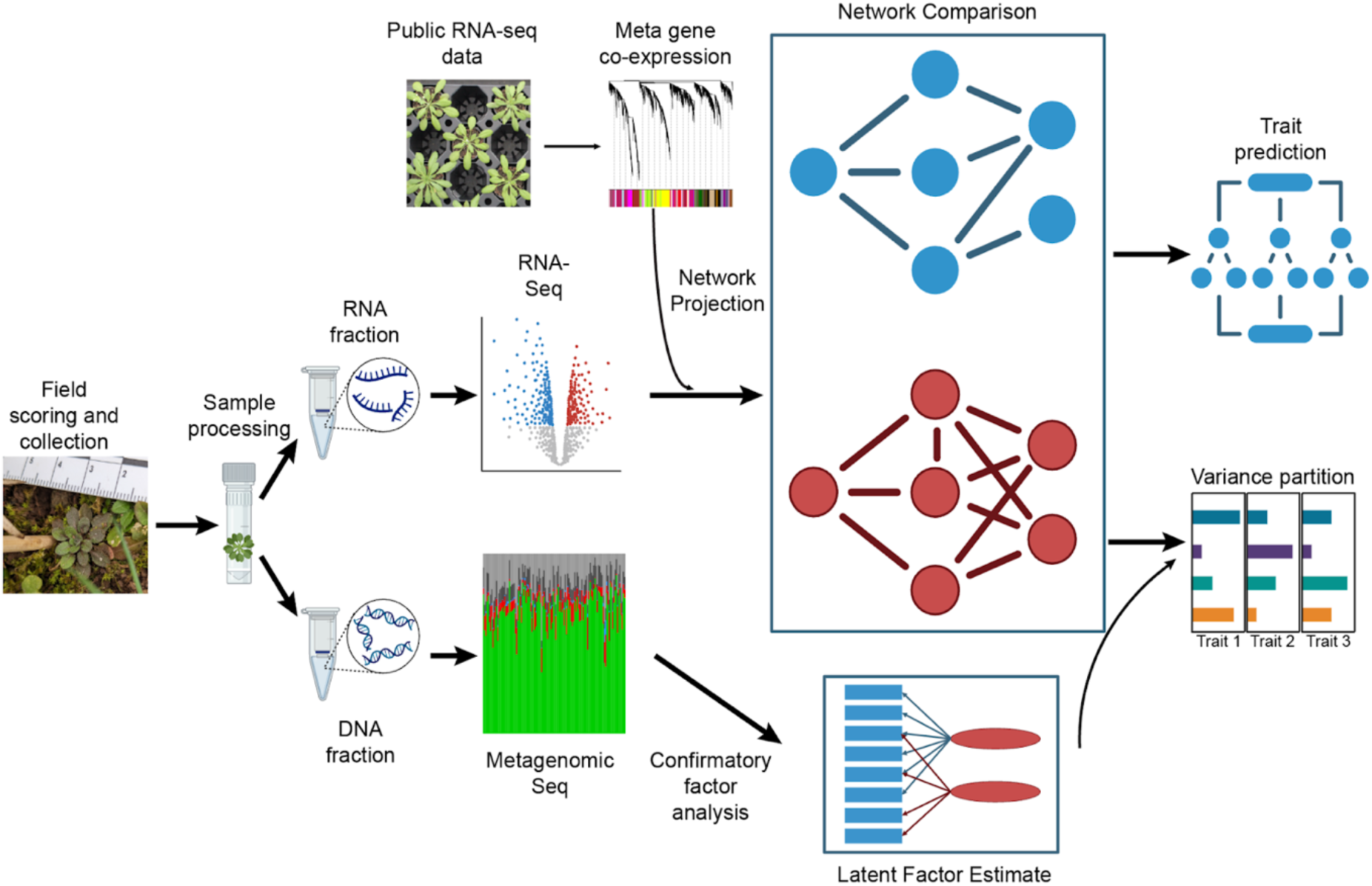

## Introduction

Over the past decades, numerous experiments in controlled laboratory conditions have revealed genetic and molecular pathways controlling development, metabolism, stress physiology and immunity. While insights from these studies have helped to improve the performance of crop plants or to develop treatments for human disease, translation from the bench to the field or bedside often fails (Des Marais *et al*. 2013; Fournier-Level *et al*. 2011; Brachi *et al*. 2010; Weinig, Stinchcombe, *et al*. 2003; Weinig, Dorn, *et al*. 2003; Loewa *et al*. 2023). This has many different reasons, but an important one is that we still have only a poor understanding of how organisms respond to the highly dynamic and complex environments that are typical for natural settings, where multiple factors tend to vary in an unpredictable manner over time and space. In the laboratory, researchers use defined perturbations on genetically identical individuals in a uniform environment, thereby maximizing the chances to identify causal effects. In contrast, organisms in the wild experience overlapping abiotic and biotic stresses throughout their life cycle, and their responses to a new challenge may depend on prior exposure and developmental state as well as other interacting species present in the environment. Understanding how co-expression networks are reorganized in such highly dynamic and complex surroundings is thus essential for bridging laboratory and field biology (Lundberg, Bergelson, *et al*. 2025).

Plants offer the advantage that they are sessile and that variation in their local environment over their life cycle is more easily recorded than for animals. Among plants, *Arabidopsis thaliana* stands out due to a wealth of laboratory studies that have probed its responses to many different environmental stressors. This in turn has led to the identification of many molecular regulators that shape these responses, but it remains difficult to predict how important these factors are under natural conditions. A prime example comes from the study of *Flowering Locus C* (*FLC*) and *FRIGIDA*, which together explains by far most of the flowering time variation among natural strains of *Arabidopsis thaliana* in the greenhouse (Lempe *et al*. 2005; Atwell *et al*. 2010), but which have much more limited effects when plants are grown in outdoors common gardens or directly in their native habitats at the time of year that matches the phenology of natural cohorts at the respective location (Brachi *et al*. 2010).

Transcriptome profiling provides a less observer-biased, high-dimensional molecular phenotype that can capture responses beyond pre-selected, visually scored traits. The study of transcriptome dynamics in the field has a considerable history, although the overall number of such studies is still limited (Nagano *et al*. 2012; Nagano *et al*. 2019; Richards *et al*. 2012; Groen *et al*. 2020; Plessis *et al*. 2015; Bhaskara *et al*. 2023; Gurung *et al*. 2019; Walter *et al*. 2023; Arana and Picó 2025; Mjema *et al*. 2026). Most outdoor transcriptomic studies deliberately reduce microenvironmental and life-history heterogeneity and thereby partially disentangle genotype from local ecology by, for example, using common gardens or sampling native plants with a design that minimizes inter-annual variation. It therefore remains unclear how transcriptome organization is structured in wild individuals, where genetic background is nested within and shaped alongside local environments and phenotypes.

To complement prior work, we examined transcriptome organization directly in wild *A. thaliana* populations across diverse sites and seasons, capturing naturally co-varying genetic, ecological, and phenotypic variation rather than controlling it. Instead of testing responses to single factors, we asked how transcriptional regulation is structured under the compound biotic and abiotic conditions typical of natural environments. Specifically, we asked (i) which environmental and phenotypic factors are reflected in transcriptome variation, and (ii) how the organization of transcriptional networks in the wild compares to laboratory-defined transcriptional programs. Although environmental and phenotypic factors explain little total variance in wild transcriptomes, infection-associated signals account for more variation than measured abiotic variables. This aligns with wild transcriptomes closely reflecting the activity of laboratory-defined biotic-response networks, whereas untreated and abiotic networks are substantially reorganized. Together, our findings suggest that while core functional modules identified in laboratory experiments remain recognizable in natural populations, their organization and activity differ, and that laboratory studies have likely only explored a small portion of all possible transcriptome states. Our study thus provides a systems-level view of how transcriptional regulation operates in natural settings and highlights the importance of integrating ecological context into molecular studies of organismal responses.

## Results

### Overall patterns of transcriptome variation in wild samples

To assay the transcriptional state of wild *Arabidopsis thaliana* plants, we sampled up to 12 plants from about 20 sites each across three study regions in Germany, France and the USA (total 60 sites, Fig. 1A-B) over two seasons (fall 2022 and spring 2023, Fig. 1D). For each plant, we measured plant traits including microbial abundance and diversity, field scores of disease, and developmental state, along with recent weather, to capture seasonal variation in factors that may shape the transcriptome (Fig. 1C-D, 2A-B; Fig. S1, S2, Table S1,S2). Since our sampling scheme necessitates visiting multiple sites per day, the time and date when plants were sampled were also recorded. Plants were washed with sterile distilled water to remove larger soil particles, and both DNA and RNA were extracted from each plant. Shotgun sequencing of DNA captured both epi- and endophytic microbial load together with plant genotype (Regalado et al. 2020; Extended Data Table 1), but for RNA sequencing data we only considered reads that mapped to the *A. thaliana* reference genome (Extended Data Table 2).

**Figure 1.**
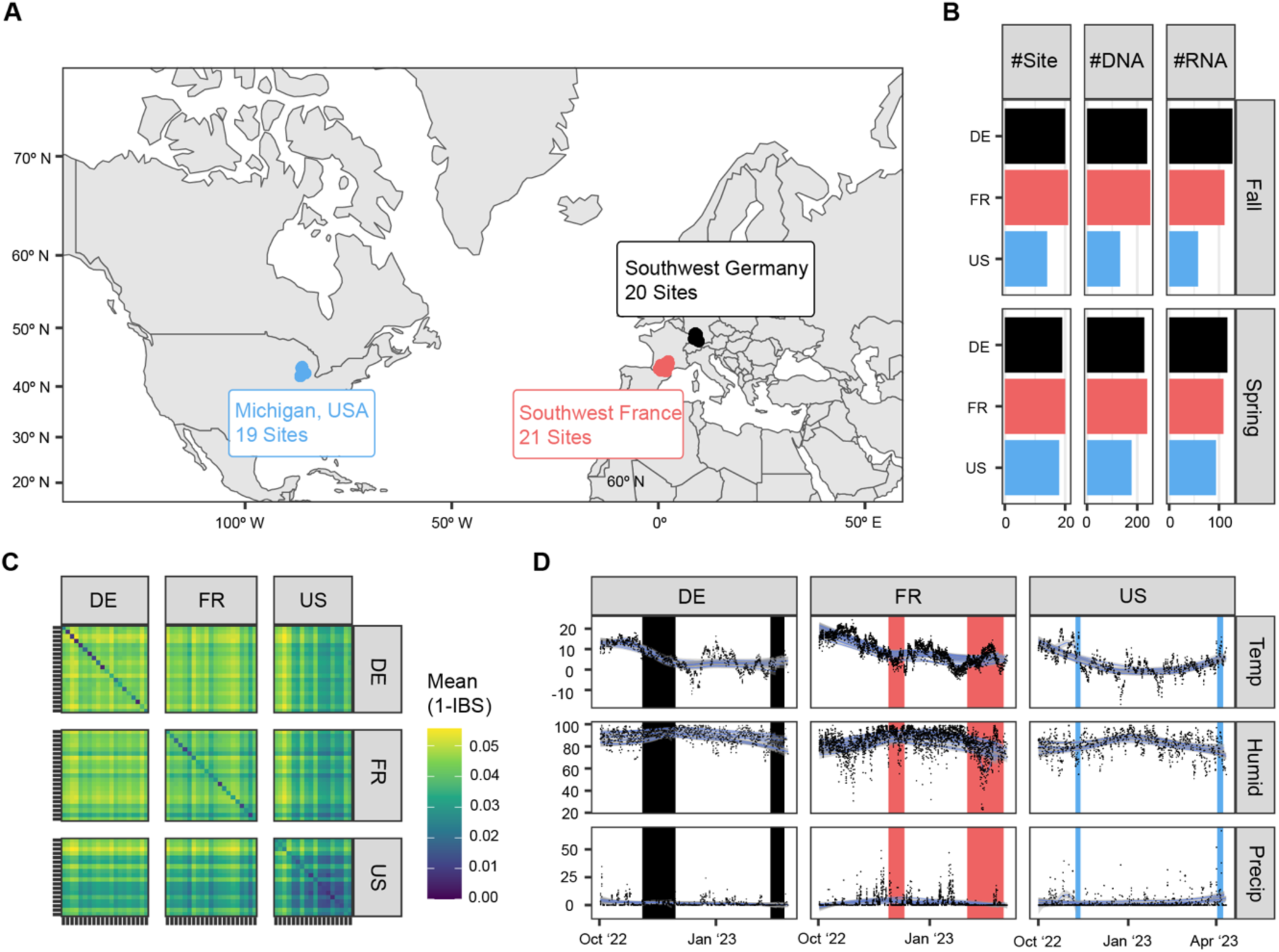
Summary of sample collection and preparation. **A,** Samples were collected from 60 sites across southwest France (FR, 21 sites in November to December 2022; 20 sites in January to March 2023), southwest Germany (DE, 20 sites in October to November 2022, 19 sites in February to March 2023), and Michigan, USA (US, 14 sites in October to November 2022, 18 sites in April 2023). **B,** Number of samples where DNA- and RNA-sequencing results passed our quality thresholds. **C**, Genetic diversity (measured as 1-mean Identity by State; IBS) within and between countries. **D**, Average daily temperature (°C), humidity (%RH), and total daily precipitation (mm) across the 2022-2023 growing season, as recorded by nearby weather stations (See Supplementary Methods). Points are days per site, and blue lines with grey bars are GAM-smoothed running means. Colored vertical bars represent the sampling periods in each country.

Of 1269 plants collected, we obtained 616 transcriptome samples, 1267 metagenome profiles, and 1092 host genotype calls, with 616 samples having all data types (Fig. 1B). The phenology of plants collected in the wild broadly aligned with the known life history of *A. thaliana* and its pathobiota (Bartoli *et al*. 2018): plants were larger, more likely to be flowering, and had more signs of disease in spring than in fall, as fall cohorts were more likely to have germinated only recently (Baskin and Baskin 1983; Donohue *et al*. 2010; Fig. S2B-D). Symptoms associated with bacterial, oomycete or fungal disease were not strongly correlated with each other (Fig. S2C-D, S3B), with the exception of chlorosis and necrosis, which are both indicators of possible bacterial infection, among other causes (Jakob *et al*. 2002; Duque-Jaramillo *et al*. 2023; Bartoli *et al*. 2018; Roux and Frachon 2022). A composite bacterial infection score was hence calculated as the sum of chlorosis and necrosis scores for each individual (Fig. 2A). Patterns of genetic relatedness within and between populations largely followed previous observations, with US populations being genetically most uniform, and French populations being most diverse (Bomblies *et al*. 2010; Shirsekar *et al*. 2021; Frachon *et al*. 2018; Fig. 1C, Fig. S1C, S2A). Microbial load and composition differed considerably among plants, even within populations (Fig. 2B; Fig. S1B).

**Figure 2.**
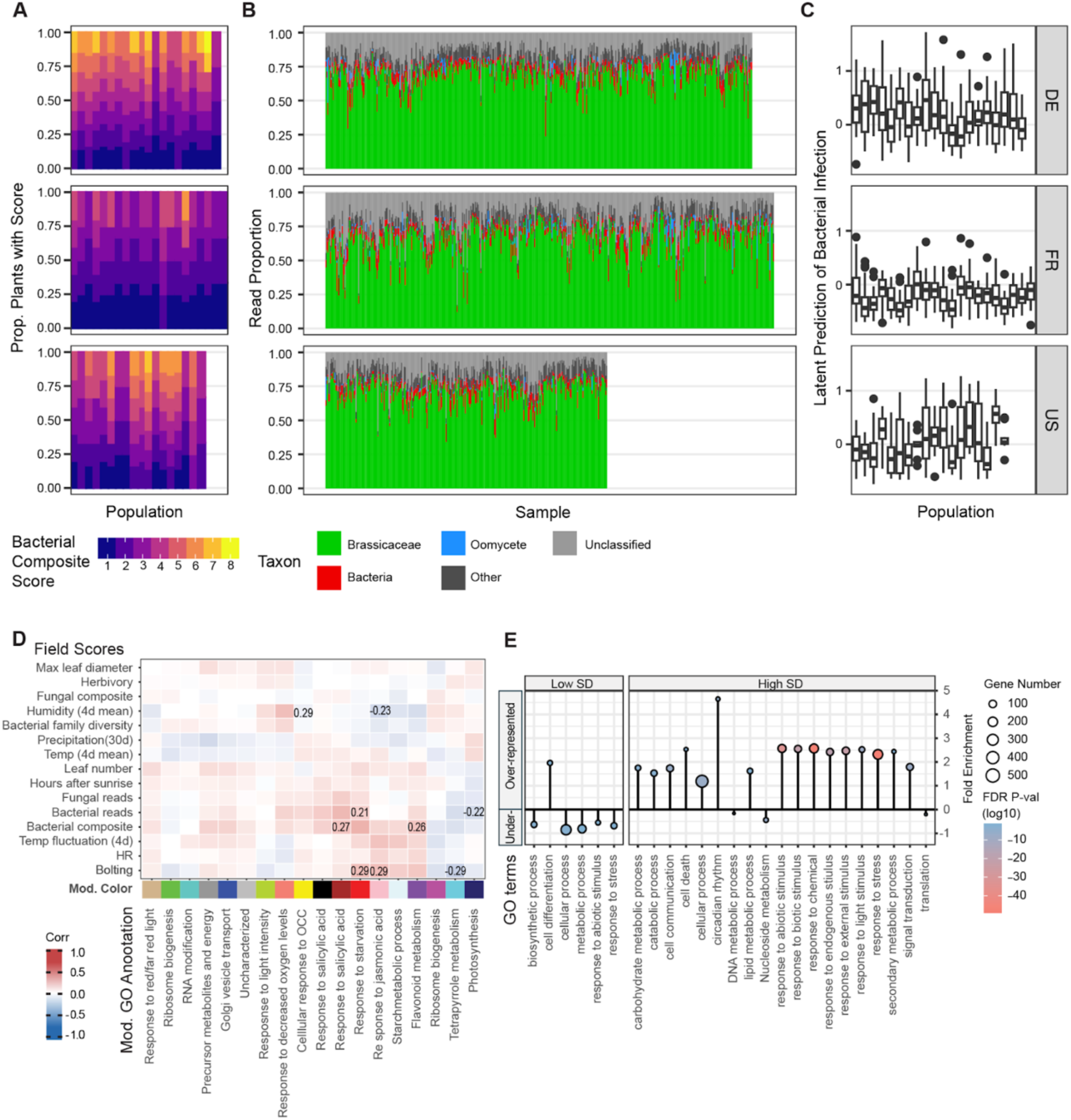
Genetic, phenotypic, and transcriptome variation. **A,** Composite bacterial scores, which are the sums of all symptoms suggestive of bacterial infection. Variation exists among both sites and countries, with the fewest symptoms in France. **B,** Microbial load and composition. Samples are ordered by site and season. The fraction of microbial reads in the DNA samples, a proxy of microbial load, varies from under 5% to 75%. *Pseudomonas*, *Sphingomonas*, *Hyaloperonospora*, and *Albugo* are the microbial taxa with the most assigned reads. **C,** Bacterial infections, as estimated with confirmatory factor analysis. **D,** Heatmap of Pearson’s r between WGCNA module eigengenes (MEs) and observed plant and environmental variables. Red indicates positive and blue negative correlation. Modules are labeled by top enriched GO biological process terms, together with the designated color labels (colored squares). Exact values are given for correlation coefficients >0.2 and p value < 0.05. **E,** Under- and over-represented GO terms for low-SD and high-SD genes. Circle size indicates number of genes in category, and color represents the FDR-corrected p-value, with red representing highly significant associations.

Variation in microbial load, defined as the ratio of DNA sequencing reads taxonomically assigned to microbial genomes versus *A. thaliana* genomes, was larger within countries than between countries, with country, population, and season explaining 3%, 19%, and 9% of variation in total non-plant load respectively (ANOVA, *p* < 1e-10). Visible signs of disease correlated variably with microbial load: correlations were relatively strong for *Albugo* spp. (*r* = 0.75; *p* < 2e-16), a major pathogen of *A. thaliana* (Taguas *et al*. 2025; Thines *et al*. 2009). Correlation was weaker for *Hyaloperonospora* spp. (*r* = 0.22, *p* < 8e-15), likely due to its longer asymptomatic phase where an infection is present but not visible. Correlation was weaker again for bacterial pathogens (*r* = 0.14, *p* < 3e-6; Fig. S3B), likely due to both asymptomatic infections (Goss and Bergelson 2007; Bartoli *et al*. 2018) and difficulties in scoring bacterial disease symptoms for small plants. We therefore performed Confirmatory Factor Analysis to synthesize latent estimates of infection from these multiple predictors of uncertain accuracy (Fig. 2C; Fig. S3).

Exploratory analyses with principal component analyses (PCA) of transcriptomes and microbiomes showed data points to be widely dispersed with no clear clusters of either transcriptome or microbiome samples (Fig. S1A-B). Compared to between-population and between-season contrasts, we had expected the median transcriptome distance to be lower within populations or geographic regions and within each season, but this was only marginally the case (Fig. 3A). Similarly, the transcriptome distances among pairs of plants with recent kinship (estimated relatedness ≥ 0.125, *i.e*., first cousin or closer) was only marginally lower than for unrelated pairs from the same population and season (Fig. 3A). These results almost certainly reflect the highly individualized life histories of even genetically closely related plants in a single wild population, with different germination dates, heterogeneous development and idiosyncratic microenvironments.

**Figure 3.**
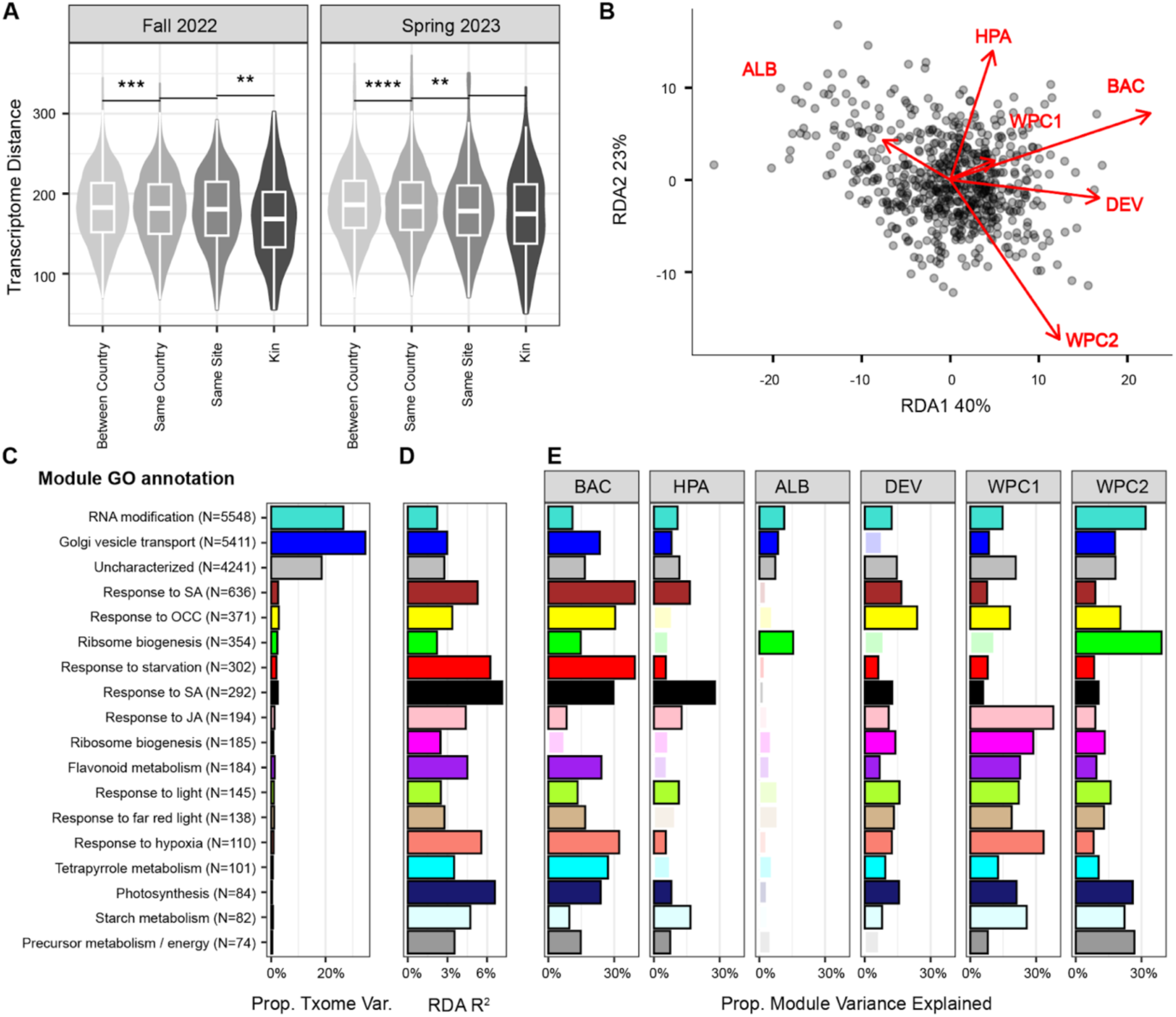
Factors explaining transcriptome variation. **A,** Distribution of pairwise transcriptome distances from individuals grouped by geographic and genetic similarity. Significant differences between pairs indicated with one or more asterisks (Wilcoxon test). The overall effect of geography or genetics on transcriptome distances is small, however as expected transcriptomes were on average slightly more similar within sites and among kin. **B,** RDA analysis of the global transcriptome, constraining transcriptome variation by latent factor predictions of infection by bacteria (BAC), *Hyaloperonspora* (HPA), and *Albugo* (ALB), and by latent factor predictions of developmental stage (DEV), and principal components 1 and 2 of recent weather obtained from nearby weather stations (WPC1 and WPC2). Overall, constrained axes explained 5% of total variance. **C,** Proportion of global transcriptome variance (Prop. Txome Var.) attributable to each of 18 wild-sample-derived WGCNA modules and number of genes in each module. **D,** Proportion of module expression variance explained in per-module RDA analyses, constrained by the same constraining variables as in **B**. **E,** Proportion of module expression variance uniquely explained by each constraining variable (see **B**). Terms with transparent colors are not statistically significant at FDR-corrected p < 0.05. Across all modules, mean proportions of constrained variance explained are 21% (BAC), 12% (HPA), 5% (ALB), 12% (DEV), 17% (WPC1) and 16% (WPC2). Colors in C-E match module color names used in Fig. 2D.

To examine the underlying biological mechanisms driving the observed transcriptomic variability, we turned to Weighted Gene Co-Expression Network Analysis (WGCNA), a powerful framework to cluster genes into modules based on their expression patterns (Langfelder and Horvath 2008). WGCNA grouped genes into 18 modules (Table S3) with correlated expression, which we functionally annotated using GO enrichment (Fig. 2D). Most modules were functionally annotated as response to stimulus, and others were associated with metabolism, photosynthesis, and biogenesis. The remainder of genes were classified into three large modules, one related to RNA modification, one to Golgi transport and one without significant GO enrichment (Table S4). The three largest groups mostly contained genes with low mean expression and low expression variance.

To determine possible drivers of correlated gene expression in each module, we asked how plant morphology, field scores and microbial load correlated with the first principal component of gene expression in each module, *i.e.* the module eigengenes (MEs), which explained 15%-45% of module variance (Fig. S4, Table S5). While most correlations were not significant, bacterial load and composite scores were positively correlated (r = 0.21 and 0.27, p-value = 0.001 and 0.025, respectively), with multiple modules being associated with response to Jasmonic Acid (JA) or Salicylic Acid (SA), and negatively correlated with the module annotated as having role in photosynthesis (r = -0.22, p-value = 0.046; Fig. 2D).

To better understand the variance distribution of the wild transcriptome, we identified the 5% most and least variable genes based on the standard deviation of normalized expression across all samples (high- and low-SD genes; Fig. S5A; Table S6). While low-SD genes were exclusively at the extreme ends of mean expression levels, high-SD genes tended to be expressed at intermediate levels, consistent with the expectation that one has the greatest power to detect variation for genes that can easily vary in both directions from their mean levels (Fig. S5B). High-SD genes were strongly enriched for GO categories related to response to various endogenous and external stimuli but depleted for house-keeping functions, while this trend was reversed for low-SD genes (Fig. 2E), in agreement with wild plants being under constant exposure to environmental stimuli. Due to the study design, we cannot rule out that expression variation in these genes can at least in part be attributed to higher genetic and/or regulatory diversity in these genes. A comparison of mean pairwise genetic distance (*π*) among all samples at each gene (±1 kb) found no significant difference between low- and high-SD genes (Wilcox Test W = 421636, p-value = 0.2374), nor between low-SD genes genes and 1,000 randomly selected genes (W = 429916, p = 0.5053). High-SD genes, though not significant (W = 456822, p-value = 0.05661), showed a greater trend towards having higher genetic diversity. Consistent with a possible role of local adaptation, most high- and low-SD genes were unique to a single region and season, especially the high-SD genes (Fig. S5C-D).

### Bacterial infection as main predictor of transcriptome variance

To quantify the extent to which measured environmental and phenotypic variables explain transcriptome variation, we performed redundancy analysis (RDA) on the entire transcriptome across all samples (Fig. 3B). We constrained transcriptome variation by latent factor estimates of plant development, latent factor estimates of bacterial, *Hyaloperonsopora*, and *Albugo* infection, and by principle components of environmental measurements (Table S7). We could explain only 5% of total transcriptome variance with these explanatory variables, and within this constrained signal, RDA1 and RDA2 accounted for 40% and 23% of the variance. Together, these results are consistent with many small, partially independent drivers shaping transcriptome differences between wild plants, and highlights that most variation is attributable to factors beyond our point-in-time observations of these plants. This contrasts with the typical laboratory expression experiment, in which the transcriptional response to a single, often strong, stimulus is observed across a set of otherwise highly similar plants.

To explore the factors that may drive expression variation within co-expressed gene modules, we performed RDA independently for each WGCNA module. Per-module models follow the global models above, with expression of all genes in a module constrained by latent factor estimates of infection with bacteria, *Hyaloperonsopora*, and *Albugo,* as well as estimates of developmental state and weather PCs. Similarly to the global transcriptome, the vast majority of variation in expression remained unaccounted for, with at most 6% of variance explained by these factors (Fig. 3C-D). Across modules, latent estimates of pathogen infection were again the largest driver of expression variation (bacteria 21%, *Hyaloperonospora* 12%, *Albugo* 5%), followed by the first two PCs of weather data (PC1 17%, PC2 16%) and latent estimates of developmental stage (12%). Of note, infection explained most variation in modules where GO term enrichment pointed to a role in immunity (Fig. 3E).

### Resemblance between wild transcriptome and biotic stress responses in the laboratory

We asked how the global organization of wild transcriptomes differ from those from the laboratory. Having established that biotic factors are important predictors of wild transcriptomic variance, we are particularly interested in how laboratory-derived abiotic and biotic networks compare when applied in the wild. Employing a comparative WGCNA framework, we first built reference co-expression networks from public RNA-seq data of laboratory experiments (Yu *et al*. 2022), then projected gene expression values from fall and spring cohorts of wild plants onto these networks to evaluate changes in transcriptome organization. We focused on a set of 1,007 biotic and 724 abiotic stress samples from 218 individual experiments (Table S8, Extended Data Table 3). We separately constructed biotic and abiotic WGCNs from the treatment samples, as well as an untreated network from 1,118 control samples. We used these networks for down-stream analyses including network projection, comparison, and random forest predictions.

The untreated, abiotic, and biotic networks contained 28, 47, and 20 modules, respectively (Fig. 4A, Extended Figure 1, Table S9-S10). The biotic network had a more uneven distribution of module sizes compared to the untreated and abiotic networks (Table S10), including one very large module with ∼40% of all genes. Across datasets, modules broadly mapped to six (biotic network) or eight (abiotic network) biological GO functions: response to stimuli, cell cycle/development, transcription/translation, photosynthesis, primary and secondary metabolism, auxin/growth, vesicle transport, and circadian rhythm (Fig. 4A, Table S11). Importantly, the three networks differed substantially in their organization, with the untreated and abiotic networks being more similar to each other than either was to the biotic network. The abiotic and untreated networks had weak overall connection strength, and both positive and negative connections were similarly common (density 0.05/0.07, mean absolute edge weight 0.04/0.05, positive edges 56%/60%, Fig. 4A). The biotic network featured stronger connectivity, and more modules were positively correlated (density 0.27, mean absolute edge weight 0.16, positive edges 86%, Fig. 4A). Both abiotic and biotic networks show a significant preference of stronger connections between modules with related GO terms (ΔfracW: +0.18/+0.2, permutation-FDR: 0.002/0.003), compared to the untreated network (permutation-FDR = 0.246).

**Figure 4.**
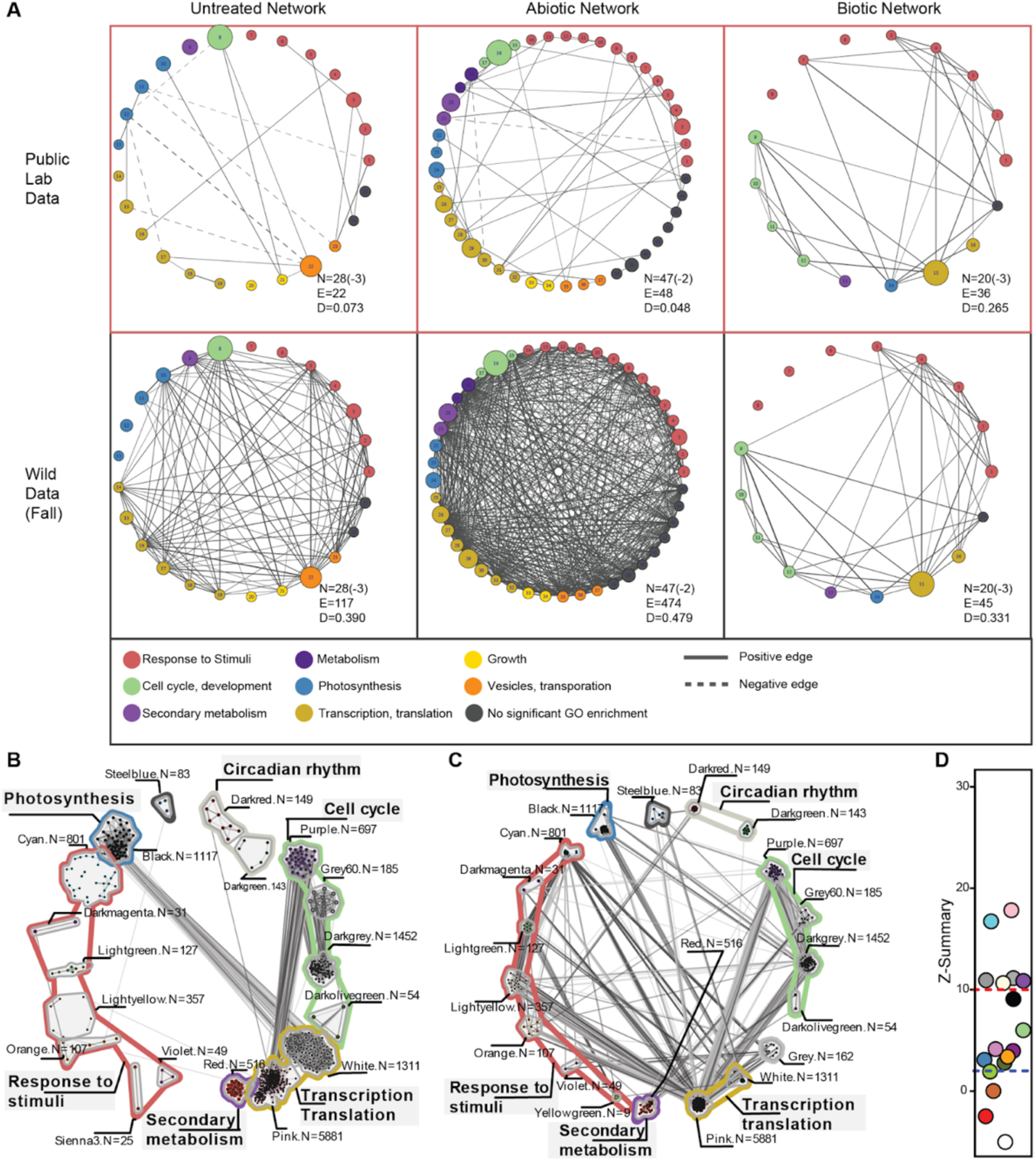
Comparative meta analyses detect conservation and divergence of modular co-expression networks. **A,** ME networks of public laboratory data (top) and of gene expression values from the wild in fall 2022 (bottom). Modules are nodes, with colors indicating GO-enrichment categories (key at the bottom). Node size indicates the number of genes in each module. Edge widths indicate strengths of connections. N: number of nodes (minus sign indicates number of circadian-rhythm-related modules); E: number of edges; D: network density. **B,** Hub-gene network from the public biotic data network (top 10% connected genes in each module). Modules are grouped by GO enrichment categories, with colored contours (color key as in A) around each group. Intensity of gray lines between genes indicates connection strength, with darker being stronger. **C,** Hub-gene network with expression values from wild samples collected in fall 2022 projected onto the biotic network. While module identities remain, hub genes are called based on the wild data. **D,** Module preservation statistics. Z-summary scores reflect the overall conservation of network topology (connectivity and density) between the laboratory and the wild. Color of dots as designated biotic WGCNA module colors (Table S11). We consider modules above the red dotted lines as highly conserved and those below the blue dotted line as not conserved. Note that modules with highest Z-summary scores are those abundant in inter-module connections in panel B.

We projected wild transcriptomes onto the laboratory-derived networks, assigning genes to modules defined by laboratory samples and estimating eigengene values for each module. Projected ME values captured on average 60% of variation in wild transcriptomes (Fig. S6), which is comparable to the proportion variance captured with the laboratory data, indicating effective dimension reduction.

Module-trait associations diverged substantially between the laboratory and the wild. Compared to the laboratory, where ME values were, as expected, strongly associated with the corresponding treatments, we did not see such strong correlations with the traits measured in the wild, regardless of whether we considered abiotic or biotic network projections (Extended Figure 2). For example, although we did observe ample indication of disease in the wild, both positive and negative associations with field disease scores were much weaker than the correlations of the same modules with microbial infections or defense hormone treatments in the laboratory. In general, the associations in wild plants were stronger in spring, which might be due to over-winter synchronization, stronger environmental cueing, or shorter life history of the newly emerged spring cohorts. Interestingly, the strong negative associations between laboratory infections and growth-related functions are most notably missing in the wild. We postulated that this might be due to an overall milder and more quantitative nature of the disease symptoms in the wild, and repeated the analysis to a subset of wild samples with the highest and the lowest disease scores. Associations in this subset are only strengthened for modules carrying response to stimuli and defense functions, but not in the growth-related modules. This suggests that while the wild plants recognize environmental assaults, the growth-defense relationship in the wild is likely modified.

Global network topology also differed substantially between the laboratory and the wild-projected networks, with the abiotic network modules becoming much more densely connected in the wild, with little apparent substructure (wild fall/spring [laboratory]: density 0.48/0.43 [0.05]; mean absolute edge weight 0.27/0.27 [0.04]; positive edges 99.7/97.0% [56%]; Fig. 4A; Extended Figure 1). A similar loss of structure was observed for the untreated network projections (wild fall/spring [laboratory]: density = 0.39/0.32 [0.07]; mean absolute edge weight 0.23/0.20 [0.05]; positive edges 93.3/93.0% [60%]).

Differently from the abiotic and untreated networks, the topology of the biotic network was strongly preserved across both seasons in the wild, with only modest increases in connectivity and density (wild fall/spring [laboratory]: density 0.33/0.32 [0.27]; mean absolute edge weight 0.21/0.19 [0.16]; positive edges 99.3/99.3% [86%]). Together, these results suggest that wild plants integrate compound abiotic stimuli over time in ways that are not reflected by laboratory treatments, which are typically either acute insults or a constant single stress. In contrast, the preserved topology of the biotic networks implies that biotic pressure may be a dominant organizer of wild transcriptomic landscapes. This seems surprising, given the mild association of modules with disease field scores (Extended Figure 2). We infer from this that either asymptomatic infections (Stergiopoulos and Gordon 2014) or lasting effects of transient infections (Lämke and Bäurle 2017) are an important factor in the wild.

We dissected how much network topology shifts were due to changes within modules versus relationships between modules. To assess changes to within-module network topology, we assayed module preservation scores (Langfelder and Horvath 2008), which separate modules into highly conserved (Z-summary > 10), moderately conserved(2 < Z-summary ≤ 10), and unconserved (Z-summary ≤ 2). Distribution of preservation classes differed significantly among networks (**χ**² = 12.79, *p* = 0.012, Fig. S7; Table S12), with a planned contrast showing the biotic networks having ∼70% lower odds to have moderately preserved modules than the others (OR = 0.30, Fisher’s Exact *p* = 0.002). Conservation also varies by function: cell cycle modules are among the most conserved, while secondary metabolism modules are among the least conserved.

To examine changes in between-module topology, we constructed hub-gene networks. Hub genes were defined as the top 10% of genes by within-module connectivity (kIN) in each module (Table S13-S14). In the laboratory biotic hub-gene network (Fig. 4B), between-module links were concentrated among cell-cycle modules, between cell-cycle and transcription-related modules, and between photosynthesis and transcription-related modules, whereas stress-response and secondary-metabolism modules remained relatively isolated. In the wild-projected biotic hub network, interconnections among cell-cycle modules persisted, but overall between-module connectivity increased markedly—most notably for hubs from the red (secondary metabolism), white (translation), and cyan (stress response) modules (Fig. 4B-C; Extended Figure 3; Table S12). A key shift was that the largest module (pink; N=5,881; GO: transcription) became a major inter-module hub, retaining its laboratory connections while gaining strong new links to multiple stress-response modules (Fig. 4C). This pattern suggests tighter coupling between growth-related regulation and stress or defense responses in wild plants via transcriptional regulation. Notably, modules showing high levels of inter-module hub gene connectivity in both the laboratory and wild are those with high conservation scores (Fig. 4D).

Abiotic hub-gene networks (Extended Figure 3) largely mirrored the ME networks (see Fig. 4A): the abiotic lab hub network showed sparse inter-module connectivity, whereas the wild-projected hub network showed pervasive inter-module connections with little evidence of module boundaries or GO-based grouping .

We also performed RDA on the wild-projected modules. Microbial infection explained more variation for modules associated with immunity or stress responses (*e.g.* module yellowgreen—systemic acquired resistance, module lightgreen— response to SA, and module cyan— response to hypoxia; Fig. S8). Taken together, these results close the loop between biotic network modules derived from well-defined, single treatments in the laboratory and our field observations. The relative lack of module association with abiotic stimuli might reflect the limited range of environments we sampled compared to the species’ native range, the severity of abiotic stimuli used in typical laboratory experiments, or failure to fully capture microenvironmental variation in abiotic factors between plants within a site and during their life history.

### Prediction of bacterial infection with module eigengenes (MEs)

To further investigate the association of the wild transcriptomes with phenotypes observed in the field, we adopted a random forest approach. We used MEs from the projected biotic modules to predict infection-related field scores. We focused on necrosis and chlorosis because of their prevalence and quantitative nature across populations and seasons. We assessed the significance of our predictions by repeating Monte Carlo cross-validation with permuted phenotype labels (1,000 permutations). Correlations with MEs were consistently higher than expected by chance (Fig. 5). Predictive performance of MEs was comparable between seasons for necrosis (median/interquartile range of Pearson’s correlation coefficients: fall 0.19/0.13, spring 0.21/0.11) and the necrosis/chlorosis composite score (fall 0.30/0.11, spring 0.35/0.10). Chlorosis alone was more predictable than necrosis in both seasons, especially in spring (spring, 0.41/0.10, fall 0.25/0.11). This suggested that chlorosis in spring might be a more integrative phenotype, for example, because it is the result of multiple disease pressures and increased vulnerability due to bolting (Extended Figure 2). In fall, the combination of necrosis and chlorosis increased predictability, suggesting partially distinct contributions of these components across seasons.

**Figure 5.**
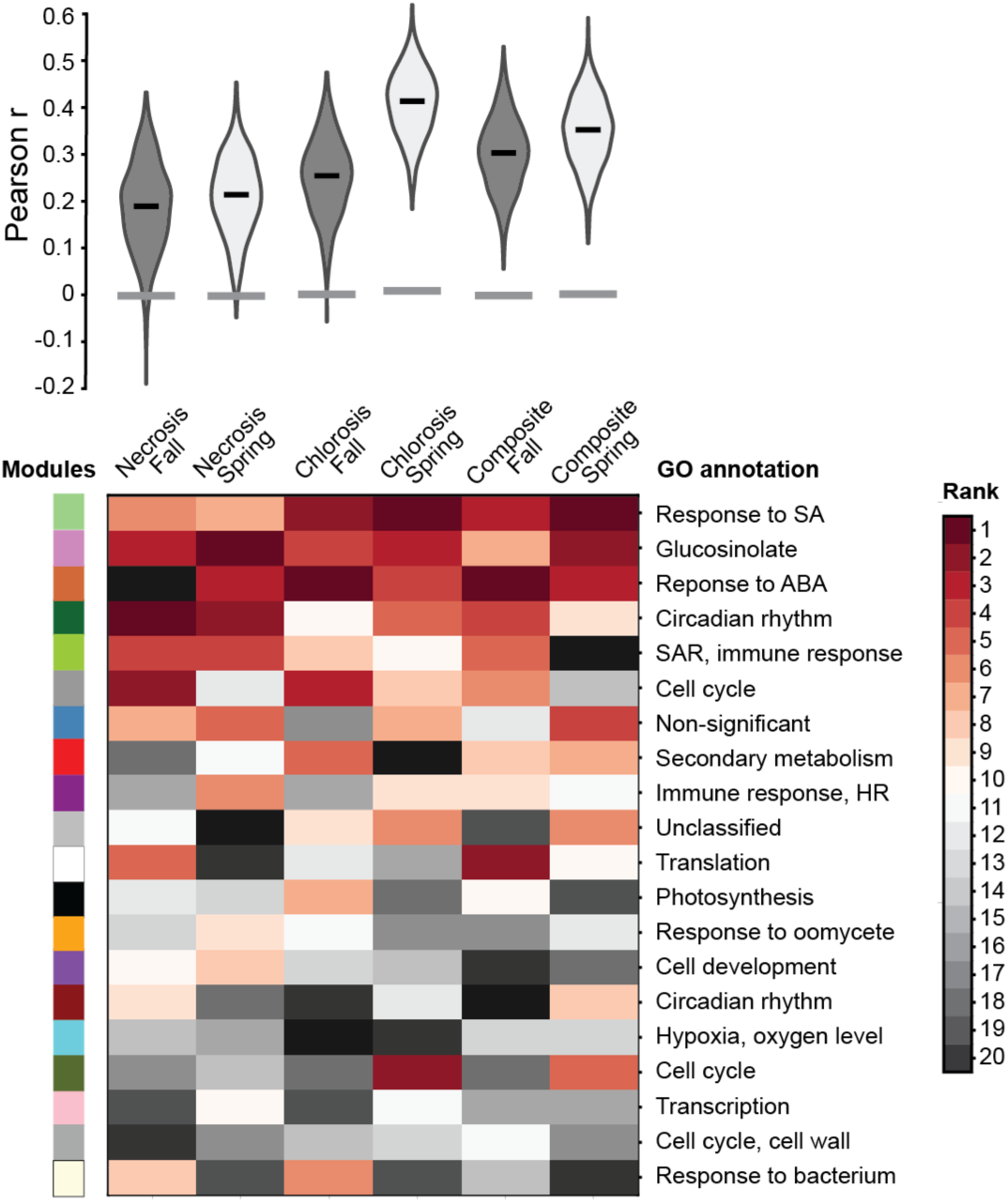
Prediction of bacterial-infection related field traits with projected MEs. Random forest prediction of necrosis, chlorosis, and their composite using MEs derived from projecting wild data onto the biotic network. Top: Pearson’s *r* between the predicted and true value. Violins illustrate the distribution of *r* in 1,000x random forest iterations. Dark grey, fall; light grey, spring. Horizontal grey bar at the bottom of each violin indicates the median value of 1,000x permuted null distributions. Bottom: Heatmap illustrating ranks of feature importance scores for each ME in the six predicted traits-season combinations. Modules are arranged from top to bottom based on their median ranks for the six combinations. Module annotated by color, see Table S11.

Across traits and seasons, most MEs (16 to 20 of 20) performed significantly better than the null expectation (Fig. S9). Five modules repeatedly ranked among the top predictors (Fig. 5). They were enriched for Salicylic Acid response (median rank 2.5, statistically significant traits 6), Abscisic Acid (ABA) response (median rank 2.5, statistically significant traits 5), glucosinolate biosynthesis (median rank 3, statistically significant traits 6), systemic acquired resistance/immune response (median rank 4.5, statistically significant traits 6), and circadian rhythm (median rank 5, statistically significant traits 6). These findings corroborated that transcriptome variation is physiologically relevant, with infection-related modules explaining a reproducible fraction of the observed disease-score variance. Modules enriched for cell cycle and translation also frequently contributed to predictions, in agreement with coordination between growth-associated and defense signaling in natural conditions.

### High variation in expression of general non-self response genes

We analyzed the expression of genes implicated in general non-self response (GNSR), which were consistently induced when *A*. *thaliana* is experimentally exposed to different bacteria (Maier *et al*. 2021). We compared the proportion of expression variance explained by latent estimates of plant state between the whole transcriptome and GNSR genes using PERMANOVA, and tested statistical significance by comparing the variance explained by 24 GNSR genes to that of 24 random genes, bootstrapped 1000 times. We found that while latent estimates explained little variance in the global transcriptome (9% total model, bootstrap range 5.13-15.23%, Fig. 6A), field scores explained a higher proportion in the GNSR genes (19% total model, p=0.001, Fig. 6A). Individually, bacterial and fungal infection, herbivory, habitat, mean temperature, total precipitation, and season had significantly higher explanatory power for GNSR genes than for random gene bootstraps (Fig. 6A). These results together confirm the GNSR genes are particularly responsive to infection by multiple pathogens.

**Figure 6.**
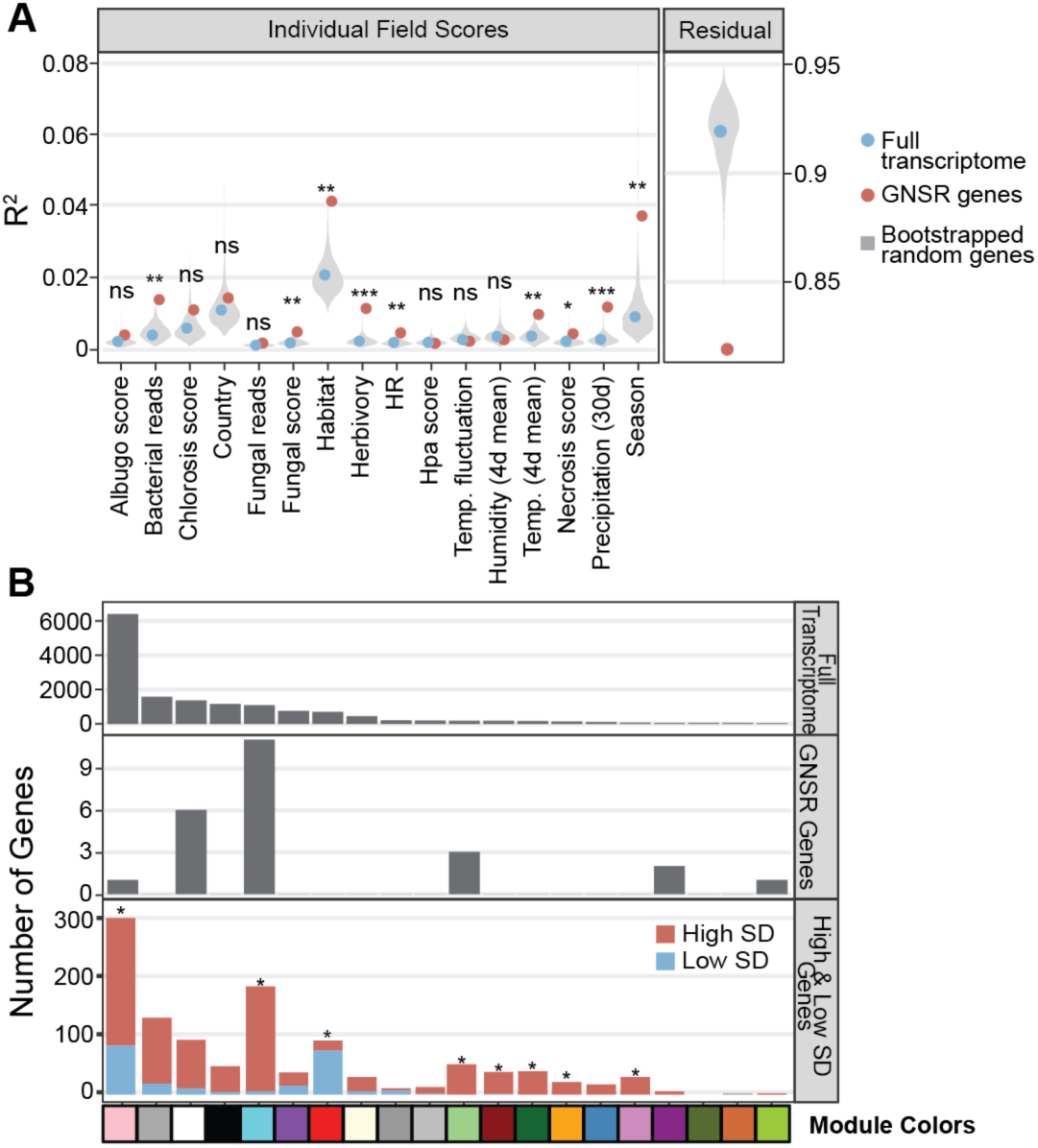
Association of general non-self response (GNSR) gene expression with field scores. **A,** Variance explained by various field scores, as inferred by PERMANOVA. Each variable was used to model variance in the full transcriptome (blue), among the 24 GNSR genes (red) and 1000x bootstraps of 24 random genes (grey violin). Asterisks indicate significance in contrast between GNSR genes and the null distribution of 24 random genes. ***, p<0.0005; **, p<0.005; *, p<0.05; ns, non-significant. Residual variance after PERMANOVA is also lower in GNSR genes in comparison to both the full transcriptome and the null samples of 24 random genes. **B,** Distribution of GNSR genes (middle) and low- and high-SD genes (bottom) does not follow the distribution of module size (top). Asterisks indicate significant enrichment of high-SD genes in a module (Fisher’s exact p<0.005). Note that most of the low-SD genes (710 of 920) were not found in WGCNA modules, in agreement with limited expression variation. For module annotation by color, see Table S11.

GNSR genes were predominantly found in biotic WGCNA modules with annotated roles in responses to stimuli (4/8 modules) and transcription/translation, and whose expression was correlated with bacterial infection (Fig. 6B, Fig. S8). We classified 13 of the 24 GNSR genes as having high expression variation (High-SD; Fig. 6B). We also found that beyond GNSR genes, high-SD genes generally were similarly overrepresented in modules with annotated roles in responses to stimuli (5/8) and transcription/translation (Fisher’s exact p-value < 0.005, Fig. 6B), which underlies that the connections between these two functions are strengthened in the wild. Together, these results suggest that responses to the multitude of challenges faced by each plant drives highly variable expression of key responsive genes, and that genes with highly variable expression, far from being simply noisy, may prove a key resource for understanding the transcriptional states of wild plants.

## Discussion

In this study, we asked how much the organization of transcriptomes into functional modules aligns between wild *Arabidopsis thaliana* plants and plants that have been grown under defined laboratory conditions. From sampling wild plants across multiple regions and seasons and combining host transcriptomes with environmental, microbial and phenotypic data, we discovered extensive transcriptome variation, which cannot be simply attributed to geography, season, growth stage, or obvious biotic or abiotic stress phenotypes. Transcriptomes were, however, not randomly organized. Using gene coexpression networks derived from laboratory datasets as a guide, we found that wild transcriptomes most closely resembled networks revealed in response to biotic challenges in the laboratory, whereas networks derived from control conditions or abiotic stresses were substantially reorganized in the wild. The absence of discrete clusters of distinct global transcriptomes in the wild suggests that the degree to which these networks are deployed is largely continuous – in stark contrast to the discrete activation of different transcriptional programs observed in response to individual stressors in the laboratory.

Redundancy analyses indicated that measured environmental and phenotypic factors explain only a small fraction of total transcriptomic variance, most likely reflecting the complexity of gene expression states in natural populations. Importantly, however, within this constrained component of variance, latent variables representing microbial infection explained more variation than developmental stage or recent weather summaries. We do not think that this implies a lesser importance of abiotic factors in nature, but rather that, under the range of environments sampled here and given the variables measured, infection-related signals leave a more detectable imprint on transcriptome variation.

Network comparisons provided additional insight into how transcriptional regulation apparently differs between laboratory and natural contexts. Modules representing core biological processes, such as transcription, metabolism, growth-related functions and stress responses, were recognizable in our data, supporting the view that functional organization of the transcriptome is broadly conserved between the laboratory and the wild. However, relationships among modules differed substantially. In particular, abiotic response and untreated networks from the laboratory lose much of their modular structure in the wild, while biotic-response networks retain much of their topology. Thus, transcriptional programs characterized under biotic challenges in the laboratory map more coherently onto expression states observed in wild populations than those derived from abiotic perturbations or untreated controls.

A notable feature of wild transcriptomes is the increased connectivity between modules associated with stress and defense responses and those linked to transcriptional and translational regulation. Hub gene analyses showed that modules related to transcription and translation occupy more central positions in projected wild networks, forming connections with both stress-response and growth-associated modules. These patterns suggest that, under natural conditions, co-expression programs associated with growth and defense are more tightly coordinated than what is observed in the laboratory, where responses are often measured after acute stimuli and in isolation.

The relationship between growth and defense responses has long been discussed in terms of tradeoffs, often, but not exclusively, based on laboratory observations where constitutive immune activation generally leads to growth penalties (van Wersch *et al*. 2016; Todesco *et al*. 2010; Mauricio 1998). Direct measurements of defense costs in natural populations or even crops have, however, yielded mixed results (Brown 2002; Bergelson and Purrington 1996; Züst and Agrawal 2017; Ojha *et al*. 2022; MacQueen *et al*. 2016; Korves and Bergelson 2004; Lundberg *et al*. 2025). Our results offer a potential systems-level perspective: differences in network connectivity between laboratory and wild conditions may contribute to differences in observed growth–defense relationships across environments. In natural populations, condition- and context-dependent regulation may reduce sustained growth penalties, allowing defense responses to operate without the strong growth suppression often observed when immunity is constitutively induced in the laboratory. Testing this possibility directly will require integrating network analyses with explicit measurements of growth and fitness in the wild.

Another striking observation is the large fraction of transcriptome variance that remains unexplained by measured variables. This likely reflects both biological and technical factors. Sampling from the wild inherently captures individuals with heterogeneous developmental histories and microenvironmental exposures that are difficult to measure comprehensively. Weather summaries derived from nearby stations may not accurately represent microclimatic variation experienced by individual plants, and we also do not know how exactly plants integrate environmental exposures over longer time scales. Soil properties may also contribute to the observed patterns. However, consistent soil measurements are not available across the full, final sample set. In addition, genetic differences among sampled individuals may contribute to variation that cannot easily be partitioned using available metadata.

Analyses of genes previously identified as part of the general non-self response (GNSR) (Maier *et al*. 2021) further support the view that immune-related pathways are dynamically regulated in natural populations. These genes show particularly high variability and stronger associations with measured field factors than expected by chance, and are concentrated within modules related to stimulus responses and transcriptional regulation. While we do not know whether this variability is indeed adaptive, it indicates that genes broadly responsive to microbial signals remain highly dynamic in wild populations and contribute substantially to observed transcriptional heterogeneity.

Our study has a series of limitations that could be addressed in future work. Disentangling the multitude of potential drivers of transcriptome variation in natural populations may require reducing heterogeneity by focusing on individuals based on synchronized germination and shared life history or measuring environmental variation at finer spatial and temporal resolution. Dense longitudinal sampling of plants across developmental stages could further clarify how transcriptome states integrate environmental history.

An informative contribution in this regard comes from a recent investigation of morphological phenotypes and transcriptomes of native *A. thaliana* plants (Mjema et al. 2026), a study that offers complementary insights to our work, based on sampling hundreds of individuals per population and year at two native sites in Germany across five years. By focusing on leaf-surface temperature measurements, Mjema and colleagues (2026) identified the expression of known key regulators of thermomorphogenesis as predictors of temperature-associated transcriptome variation. In comparison, our broader geographic sampling, which came at the cost of the number of samples per site, shows that, once the seasonal contrast is set aside, the dominant organizer of transcriptome variation is biotic in nature, with infection-associated signals accounting for more of the explainable variance than recent weather summaries. We conclude that temperature and biotic stress are not competing explanations for the organization of wild transcriptomes, but rather operate at different temporal and spatial scales. Temperature impacts broad seasonal and year-to-year trajectories detectable when the same populations are sampled at depth, whereas pathogen-associated signals emerge as the primary structuring force when transcriptomes are sampled across the full ecological heterogeneity of the species’ range. Finally, while most of the variation in native transcriptomes remains insufficiently explained by variables measured in the field in both studies, together they begin to provide a path forward for designing sampling strategies across space and time that capture more of the factors that shape native transcriptomes.

At a broader level, our results support a systems-level perspective in which functional gene modules identified in laboratory experiments remain relevant in natural populations, but their organization and activity differ across environments. Laboratory experiments excel at identifying modules and causal pathways under defined perturbations, whereas natural environments reveal how these modules are coordinated under sustained and heterogeneous conditions. Integrating both approaches offers a path toward understanding how molecular regulation operates in realistic ecological contexts and how co-expression networks are deployed across environments.

## Methods

### Wild plant sampling

Full rosettes were collected from 60 wild populations of *Arabidopsis thaliana* across southern France, midwestern USA, and southwest Germany. Sampling was conducted during fall 2022 (November-December) and spring 2023 (February-April), with exact timing depending on the local conditions in each country (Fig. 1). At each site and season, we aimed to collect 12 individual plants within two to three weeks. For each plant we recorded signs of herbivory (0-5), hypersensitive response-like symptoms (P/A), as well as symptoms of bacterial disease (chlorosis, necrosis; 0-5), which were used both individually and as a composite score of the their sum, and symptoms of infection by fungi, *Hyaloperonospora* spp., and *Albugo* spp. (0-5). (Fig. 2A, Fig. S2, Table S1). The developmental state was characterized by counting the number of leaves, rosette diameter, and the presence or absence of bolting. We extracted weather data from nearby weather stations (Table S2) from www.wunderground.com for the duration of the sampling season (October 1, 2022, to April 30, 2023). For each sampling visit, we summarised the weather data as mean temperature of the four days preceding sampling, mean humidity for the four days before sampling, mean diurnal temperature range for the four days before sampling, and sum of precipitation during the 30 days preceding sampling. We summarised these variables through the first two principal components (PCs) in a principal component analysis (PCA).

### RNA and DNA extraction

DNA and RNA were simultaneously extracted from individual rosettes. After scoring, wild-collected rosettes were washed with sterile distilled water to remove surface soil, then placed in 2-ml screw-cap tubes containing a mix of glass and ceramic beads (equal volumes of sterile 0.2mm, 0.5mm, 1.5mm ceramic beads, and one 5mm glass bead, dispensed with a Custom Lab Institute 96 well Powder Dispenser), frozen on-site in dry ice and stored at −80°C until extraction. Samples were homogenized to a fine powder using a Precellys Tissue Homogenizer (Bertin Technologies, Montigny-le-Bretonneux, France) equipped with a Cryolys cooling adapter, using two cycles of 15 s at 6,300 rpm. Nucleic acids were purified using ECONOSPIN columns (Epoch Life Science, Inc., Fort Bend, TX, USA) adapted to 96-well plates following the protocol described in (Yaffe *et al*. 2012). Briefly, 800 µl of extraction buffer (20 mM EDTA, 8M guanidine HCl, 20 mM MES) was added to each sample. Homogenized samples were mixed with equal volume of absolute ethanol before loading the mixture onto column plates. Samples were sequentially washed with 3M sodium acetate, twice with 70% ethanol before elution in 60 µL nuclease free water. Aliquots of eluates were subjected to RNase A or DNaseI (Thermo Fisher) treatments to retrieve DNA and RNA fractions, respectively. The products were stored at −80 °C.

### Library preparation and sequencing

We constructed whole-genome shotgun DNA sequencing libraries using an in-house tagmentation protocol similar to Nextera Flex, originally derived from (Picelli *et al*. 2014). Briefly, we normalised DNA to a maximum concentration of 5 ng/µL, tagmented the DNA with bead-bound Tn5 transposase, then amplified with dual-index primer pairs with 12 cycles of PCR using Q5 polymerase (NEB). Cleaned up libraries were pooled at equal volume across a plate. Plate pools were subjected to size selection (300-750 bp) using BluePippin (Agilent Technologies, Santa Clara, CA, USA), with final fragment size assayed on the Bioanalyzer platform (Agilent Technologies), and pooled across plates equimolarly. We sequenced libraries to a mean yield of 6 Gb on an Illumina Novaseq X Plus 25B instrument with 2×150bp paired end reads.

We constructed RNA-seq libraries based on an in-house protocol (Yuan *et al*. 2023). Briefly, we used ∼1 µg of total RNA from full rosettes, synthesized cDNA, selected mRNA with oligo-dT bead-pulldown, then produced libraries with unique dual-indexed barcodes. We pooled libraries at equimolar ratios, and sequenced the pool to a mean yield of 2 Gb per library.

### Metagenome analysis

We used shotgun DNA sequencing reads to determine the ratio of microbial and plant genomes, to call plant genotypes, and to assign microbial reads to different taxa. We analysed WGS reads with the Acanthophis pipeline (Murray *et al*. 2024). Briefly, we performed read QC with AdapterRemoval (Schubert *et al*. 2016), trimming adapters and low quality sequences from reads, and removing reads that were shorter than 64bp after trimming. We mapped reads to the *A. thaliana* TAIR10 reference genome (Lamesch *et al*. 2012) with BWA MEM (Li 2013), and called variants with DeepVariant and GLNexus, using the Deepvariant-WGS model (Yun *et al*. 2021; Poplin *et al*. 2018). We filtered variants as follows: samples were removed if they had less than 10 total depth (aggregated over SNP calls), or had more than 5% missing genotype calls, and sites were removed if they had more than 5% missing data, or fewer than 3 observations of the less common allele.

To allow deeper taxonomic profiles without excessive RAM use, we used a hierarchical reference set to create a custom kraken2 database. Specifically, we took all RefSeq representative bacterial genomes, and all RefSeq representative genomes from Brassicaceae. To this, we added all complete *Arabidopsis* spp. genomes from the NCBI genomes portal, and additional 77 *A. thaliana* long read genomes (Wlodzimierz *et al*. 2023; Teasdale *et al*. 2025), as well all complete genomes for every bacterial genus in At-LSPHERE (Bai *et al*. 2015). We used kraken2 (Wood *et al*. 2019) to assign all reads passing QC to this custom *A. thaliana*-specific reference database, and to the Kraken2 PlusPFP reference database. We used the proportion of basepairs assigned to each microbial family divided by either the total library yield (prop_taxon) or the number of basepairs assigned to Brassicaceae (taxon_per_plant) respectively, using R and DuckDB. We also summarised to higher taxonomic levels: Oomycota, Fungi, and Bacteria, reported as both proportions and host ratios as for family-level summaries above, along with Other, representing reads that were positively classified but not to one of these higher level groups, and Unclassified reads that could not be assigned to any taxon (Extended Data Table 1).

### Latent variables estimates of infection

To better integrate the multiple unreliable indicators of microbial infection (including visual scores and taxonomically-ambiguous microbial reads), we used confirmatory factor analysis to predict latent variables that represent the consistent signal of these indicators. To do so, we modeled infection as a function of all observed variables likely to be correlated with a genuine infection using cfa from the R package lavaan (Rosseel 2012). We modeled infections of bacteria (infection ∼ necrosis + chlorosis + hr + prop_pseudo + prop_xantho + prop_sphingo + bact_family_entropy), *Albugo* spp. (infection =∼ albugo + prop_albugo), and *Hyaloperonospora* spp. (infection =∼ hyalo + prop_hyalo), producing predictions of infection that incorporate variation across both field observations and sequencing data. We also fit a model of developmental state, as (dev_state =∼ rosette_diam_mm + num_leaves). Bolting was not included in the developmental state estimates as it is a binary trait largely nested within season leading to unstable model fit. Overall, we found the model fit to be good: cfi=0.896; tli=0.848; rmsea=0.063; srmr=0.047 (Fig. S3).

### RNA-seq analysis

We first quality controlled short RNA-seq reads with AdapterRemoval (Schubert *et al*. 2016), then down-sampled 132 high-coverage samples to no more than 10 million reads using seqtk (Li 2008). We then mapped RNA-seq reads to the *A. thaliana* TAIR10 reference genome with Araport11 annotation (Cheng *et al*. 2017), and estimated expression of individual gene models with RSEM (Li and Dewey 2011). A snakemake pipeline for these steps is available at https://github.com/kdm9/montalvao-rnaseq (Köster and Rahmann 2012). We filtered RNA count tables as follows: we removed organellar transcripts, transcripts mapped to rDNA, rRNA, nucleolar genes, pseudogenes, pre-tRNAs, and transposable elements, as well as transcripts with RSEM-estimated effective length less than 150 bp, whose count estimate tends to be inaccurate (Conesa *et al*. 2016). This left 30,669 genes for further consideration.

We then removed samples with fewer than 2 million mapped reads, leaving a total of 616 usable samples. We applied the Variance-stabilizing transformation (VST) on the filtered raw-counts of these samples using DESeq2 command vst (Love *et al*. 2014). We further filtered the normalized count matrix to remove genes with zero expression in more than 580 (95%) samples, extremely low average expression (vst≤4.2), and either extremely low (coefficient of variation ≤0.01) or extremely high variance (coefficient of variation ≥ 0.23). We determined all thresholds manually after examination of the histograms of the respective parameters. We then created the normalized expression matrix of 18,452 genes and 616 samples (Extended Data Table 2).

A first look at transcriptome variance (PERMANOVA, Dixon 2003) identified sampling time (hours after sunrise), country, and season to be the major source of systematic biological variation, while plate effects (i.e, on which 96-well plate a library had been synthesized) drives major batch noise. To correct for batch effect while preserving biologically meaningful signal, we used limma’s removeBatchEffect (Smyth 2005; Ritchie *et al*. 2015), while specifying effects to be kept using optional argument design= model.matrix(∼ hours_after_sunrise + country + season). PCA was run before and after to evaluate the efficiency of batch effect removal.

### Analysis of high- and low-SD genes

We calculated the standard deviation across samples for each gene using variance stabilized expression values. The 5% of genes with the highest and lowest SD were considered high- and low-SD genes (N=922 genes for each group). GO term enrichment was performed using PANTHER (Mi *et al*. 2019) with FDR-corrected p-values. To determine whether high- and low-SD genes were similar across countries and seasons, we subset samples (*e.g.*, France + Spring), and re-calculated standard deviation.

### De novo WGCNA generation

Signed WGCNA networks (Langfelder and Horvath 2008) were generated using biweight midcorrelation (bicor; maxPOutliers = 0.1). Soft-thresholding powers from 1-30 were evaluated using the scale-free topology criterion, and a power of 6 was selected as the lowest value achieving signed R² ≥ 0.8. Modules were identified using blockwiseModules (minimum module size = 30, deepSplit = 4, pamRespectsDendro = FALSE), and similar modules were merged at a cut height of 0.15. MEs were derived as values of the first PC of each module. Associations between MEs and external variables were assessed using Pearson’s correlation.

### Modeling transcriptome variation using latent estimates

We used Redundancy Analysis (RDA) to model variation among genes in each of the 18 wild WGCNA modules as a function of latent estimates of infection, development, and summaries of weather data. For each module, we fit an RDA across all genes in the module (removing between 0 and 2 genes per module with perfectly correlated gene expression to avoid ill-defined models). We used latent estimates of infection with bacteria, *Hyaloperonspora* spp., and *Albugo* spp. as well as developmental stage, and recent weather PC1 and PC2 as constraining factors in each RDA model. We summarised the variance in each model hierarchically as follows: proportion of total transcriptome variance contained in this module, proportion of module variance constrained by the explanatory factors, and proportion of constrained variance attributable uniquely to each constraining factor.

### Curating public RNA-seq data

We downloaded FPKM values for an ensemble of 28,164 *A. thaliana* RNA-seq samples and their corresponding metadata from https://plantrnadb.com/ (Yu *et al*. 2022). Almost all samples were from the Col-0 reference strain (Table S8). We manually curated metadata to include only experiments with more than 5 samples with either biotic and/or abiotic treatments, and removed any samples with combinatorial treatments. To enable further WGCNA module-trait associations, we manually coded a binary design matrix of treatment × sample to across 24 distinct biotic and abiotic treatment categories, assigning 1,118 control/mock samples, 724 abiotic treatment samples, and 1,007 biotic treatment samples (Table S8, Extended Data Table 3).

### Comparative meta-WGCNA

#### Curating public gene expression data

We removed genes with >50% dropout rate, mitochondrial, chloroplast, rDNA, rRNA, and nucleolar genes, pseudogenes, pre-tRNAs, transposable elements, and genes shorter than 150 bp. Genes with extremely low average expression, and globally invariant genes were filtered separately for the abiotic/biotic treatment datasets and the untreated samples based on quantile distribution (abiotio/biotic: mean log(FPKM) ≤1, SD≤1.15; untreated: mean log(FPKM) ≤1, SD≤0.99). This left 14,854 genes for the abiotic/biotic samples, and 13,630 genes for the untreated samples.

#### WGCNA network construction

We generated signed WGCNA networks (Langfelder and Horvath 2008) biweight midcorrelation. Soft thresholding power (Untreated=13, Abio=12, Bio=20) was evaluated with the pickSoftThreshold function (signed, bicor) on a series of thresholds from 1-30, and determined using scale-free fit (R² ≥ 0.8) and mean connectivity (Untreated:51.39, Abio: 63.99, Bio: 24.92). Deep split values (Untreated: 3, Abio: 4, Bio: 2) were chosen by running 100x boot straps of 80% of the samples on allowed deepSplit values (2-4) for each dataset, and evaluating the module robustness using adjusted Rand Index (ARI) and Jaccard Index. Similarly, optimal tree-cut height (Untreated: 0.15, Abio: 0.2, Bio: 0.1) was determined by running a series of cut heights, and bootstrapping the merged modules for ARI and Jaccard index as stability measurements.

We constructed gene co-expression networks with WGCNA step-wise using functions adjacency (signed, bicor), TOMsimilarity, cutreeDynamic (minimum module size=30), and mergeCloseModules. After each gene had been assigned to a module, we calculated MEs as the first PC for all genes in a module, using the WGCNA function moduleEigengenes. GO biological process enrichment was performed for each module using packages clusterProfiler (*v4.10.1*, Yu 2024) and org.At.tair.db (*v. 4.5*, Carlson 2024). GO enrichment results were manually curated to determine a single representative GO annotation for each module (Table S11). We calculated Pearson’s correlation coefficients to establish associations between MEs and treatments.

#### Network projection

For each season, genes in the wild RNA-seq data set were assigned to modules in the untreated, abiotic, and biotic networks. Expression of most genes was detected in both laboratory and wild data: of 14,854 genes included in the laboratory abiotic/biotic networks, 13,256 were also in the wild data, and for the untreated network, of 13,630 genes, 12,093 were also in the wild data. ME values in the projected networks were calculated using the WGCNA function moduleEigengenes. We calculated Pearson’s correlation coefficients to establish associations between MEs and the field scores.

#### Constructing ME networks

We calculated the ME adjacency matrices by arbitrarily raising the ME correlation matrices to the power of 4. A range of powers was tested to make sure that results with a power of 4 were representative. Adjacency matrices were then converted into edge lists, and visualized with igraph (*v 2.1.4*, Csárdi *et al*. 2026). To facilitate visualization, the edgelist was trimmed using an arbitrary hard threshold of 0.25, after testing a range of hard thresholds for representativeness. The circadian rhythm modules were excluded from the MEnetworks lest they drive major network structural changes that were simply due to different sampling times in the field.

#### Evaluation of the ME networks

To describe and compare ME network properties, we computed the number of edges (E) and network density (D; fraction of all possible module–module pairs connected), along with mean absolute edge weight |w|, and the percentage positive edges (%+E), where edge weights were signed adjacency values derived from pairwise ME correlations. To focus on the most informative connections and match network visualizations, E and D were calculated on a thresholded edges (|w|>0.25). whereas, |w| and %+E were calculated on the untrimmed signed adjacency matrix to summarize overall coupling independent of the hard-threshold choice. Functional modularity of the network was evaluated using GO assortativity (ΔfracW), defined as the observed fraction of strong-edge absolute weight occurring within GO categories minus the mean of this quantity under a null generated by permuting GO category labels across modules while holding network topology and edge weights fixed. Significance was evaluated using a one-sided permutation test for enrichment and Benjamini–Hochberg correction across networks.

#### Module preservation analysis

Topological preservation of each module was formally calculated by building a multi-expression dataset using common genes within each module between lab and wild expression data, and running the modulePreservation function with the lab network set as reference, and 200 rounds of permutations. Z-summary statistics were used to evaluate preservation status of each module.

### Hub-gene network analysis

#### Hub gene identification

Hub genes were defined as genes with the highest 10% of within-module connectivity (kIN) in either the laboratory or wild-projected networks. kIN was calculated as the sum of adjacency values only within the module, and total connectivity (kTOT) was calculated as the row-sums of the adjacency matrices. Between-module connectivity was calculated as the difference between kTOT and kIN. Hub gene sets were obtained separately for each reference network (untreated, abiotic, biotic) and, where applicable, for wild-projected networks (spring and fall) using the same module assignments as for the laboratory networks.

#### Hub-gene network construction

Networks were generated to focus on the strongest connectivity relationships among intra-modular and inter-modular hub genes. For each network, the corresponding adjacency matrix was first subset to include only the entries between hub genes. To sparsify the graph while retaining strong connections, adjacency values in the sub-matrix were subjected to an upper quantile cutoff of 0.95. Self-edges were removed and undirected edge lists were constructed from the remaining non-zero entries. Reverse duplicates arising from the adjacency matrix symmetry were removed by sorting node pairs and discarding duplicated pairs. Graph objects using the package network *(v1.19.0,* Butts 2008) were built from the filtered undirected edge list. The nodes were annotated with module membership for each gene based on the module–gene association tables and the edges with weights corresponding to the retained adjacency values. For visualization, we used the packages intergraph (*v2.0-4*, Bojanowski 2023), GGally *(v2.4.0,* Cook *et al*. 2025) and ggplot2 *(v4.0.1*, Wickham 2016). Edge colors were discretized into three bins spanning the observed weight range (light to dark grey). Modules were sorted and colored by manually curated GO categories (Table S11) and arranged on a circle. Each module’s hub genes were visualized around a position on the circle using a force-directed layout. Two levels of labelled convex hull annotations were added using ggplot’s geom_mark_hall – colored hulls grouping modules by GO category and grey hulls around individual modules.

### Random forest prediction of field scores

We used random forest regression models from the scikit-learn Python package *(v1.3.2,* Pedregosa *et al*. 2011) to evaluate the predictive ability of MEs for phenotypical or metagenomic traits.

Feature normalization was applied in the first step using QuantileTransformer from the scikit-learn package. Uniform distribution and quantile number = 100 were used as parameters. For split season experiments, normalizations were applied separately per season. Pearson correlation (*r)* and R2 score were used as measures of predictability. Permutation feature importance was used as a measure of the contribution for individual MEs. Monte Carlo cross-validation with a random 70% train/30% test split with 1,000 iterations was performed to generate distributions of *r*, R2 scores and feature importances. Null distributions were created by randomly shuffling the trait values in each train/test split, then training and evaluating a random forest regression model on these datasets.

Mann-Whitney U tests (one-sided, “greater”) were used to evaluate statistical significance, with Benjamini/Hochberg adjustment for multiple tests.

### Analysis of general non-self response (GNSR) genes

Hellinger-transformed expression values were subset to contain only the 24 GNSR genes (Maier *et al*. 2021). Euclidean distance was calculated including either the full gene list, only the 24 GNSR genes, or 1,000 bootstraps of 24 randomly chosen genes. Variance explained by different factors was estimated using PERMANOVA (adonis2), where R² represents the fraction of total multivariate variance in transcriptomic distances attributable uniquely to that factor. Significance was determined via 999 permutations.

All analyses were performed in the R programming environment (*v4.4.0*, R Core Team 2024), unless otherwise specified.

## Data availability

Sequencing reads are available at the ENA under the project number PRJEB109258. Extended data files are available at figshare at doi:10.6084/m9.figshare.31638136.

## Code availability

Analysis code is available at https://github.com/kdm9/montalvao-rnaseq/.

## Acknowledgements

We thank Luisa Teasdale for comments on the manuscript, Rebecca Schwab for help with identifying sampling sites, Patience Chatukuta, Adrián Contreras, Eirik Lågeide, Caterina Lino, Miriam Lucke, Andrea Movilli, Gal Ofir, Sheila Roitman, Katerina Romanova, Rebecca Schwab, Luisa Teasdale, and Shanshan Wang for help with sampling, and Dafne Ibarra for helping to coordinate sequencing.

## Author contributions

FR, JB and DW conceptualized the study; AM, KM, WY and DW conceived the project and analytical framework; AM, KM, NB, LH, PD and MK coordinated and performed field collections; AM, KM, NB and WY generated the sequencing data; AM, KM, IB, LH, PD, PB and WY analyzed the data; AM, KM, WY and DW wrote the first draft of the manuscript. All authors edited the manuscript. TEAM PATHOCOM assisted with field collections and lab work and contributed advice.

## Extended data

**Extended Data Table 1:** Wild metagenomic profile

**Extended Data Table 2:** Normalized wild RNA-seq read counts

**Extended Data Table 3:** Filtered logFPKM of public RNA-seq data

**Extended Figure 1:**
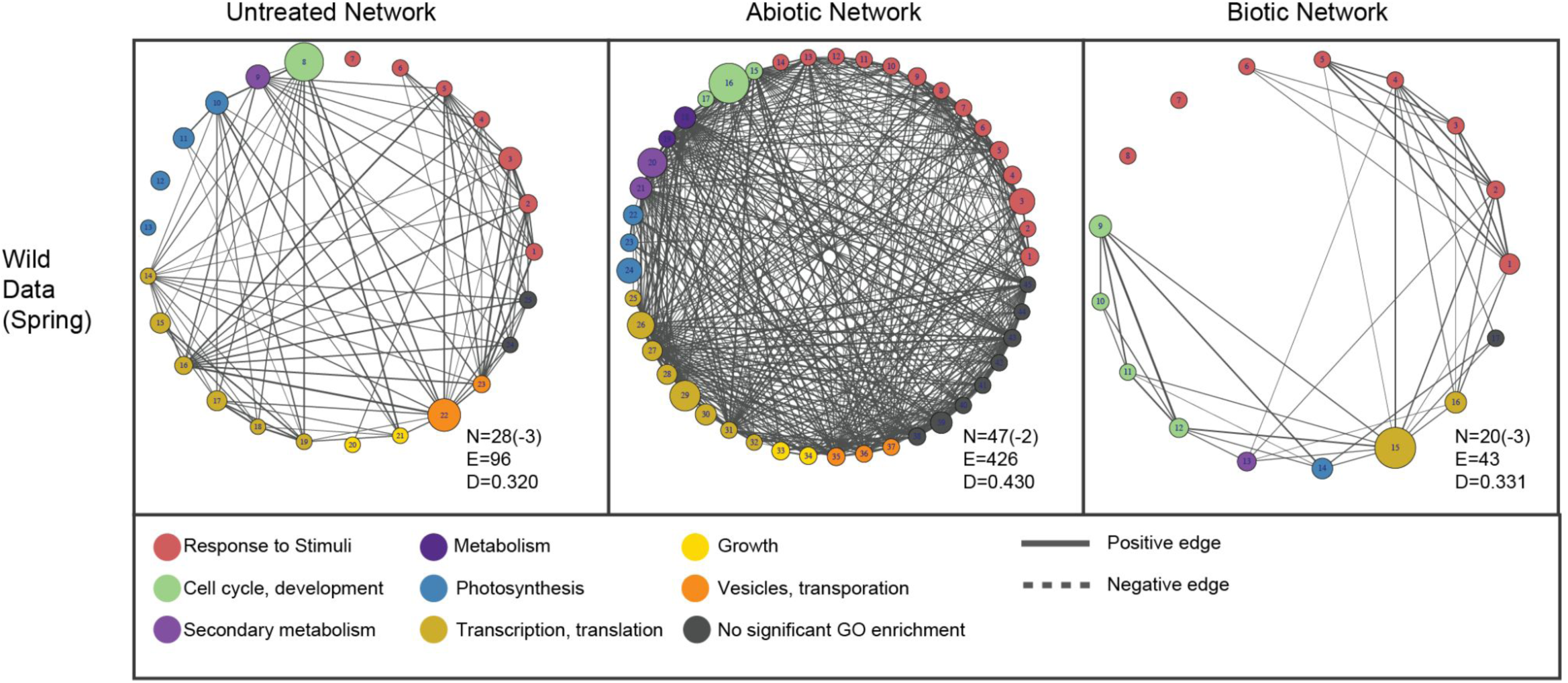
Module eigengene (ME) networks of public lab modules projected to wild samples collected in spring 2023. Each filled bubble represents a module, colored: GO-enrichment categories, size: module size, edge width: strength of connection. N: Number of nodes/modules (minus circadian modules). E: edges. D: network density. Consistent with Fig. 4A, while biotic ME network remain largely conserved between lab and projected wild data, the topology of the untreated and abiotic ME networks are heavily disrupted.

**Extended Figure 2.1:**
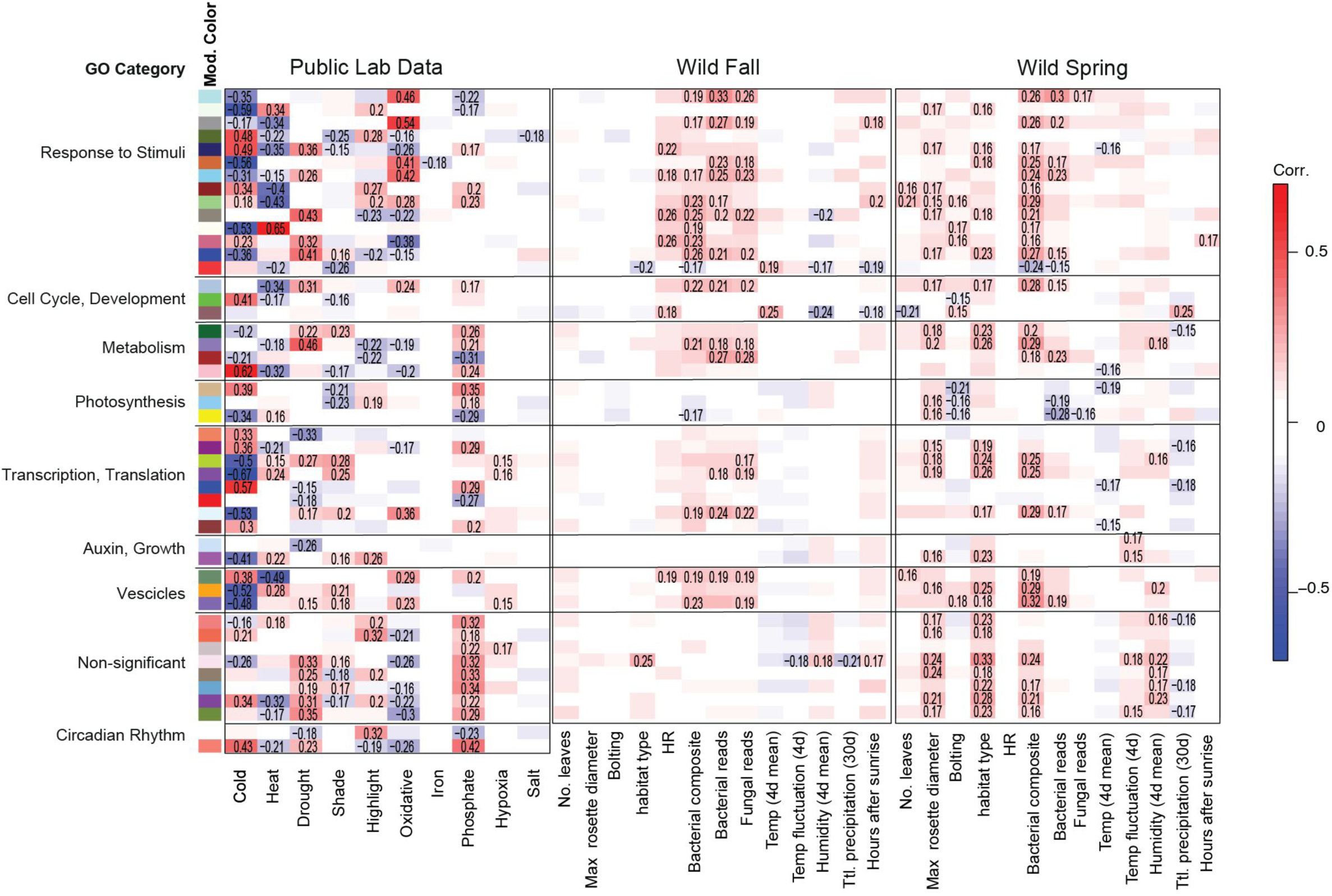
Module–trait correlation heatmaps. Pearson correlations between eigengenes (MEs) of the public abiotic WGCNA and laboratory treatments (left), and between eigengenes of wild fall (middle) and spring (right) transcriptomes projected onto the abiotic modules and their corresponding field scores. Module colors are indicated by rectangles on the left, and modules are grouped into eight GO categories. Numbers in the heatmaps show Pearson’s *r* for significant ME–trait associations only (*p* < 0.005). Lab MEs exhibit strong, bidirectional correlations with applied treatments, whereas associations between projected wild MEs and field scores are weaker but generally positive.

**Extended Figure 2.2:**
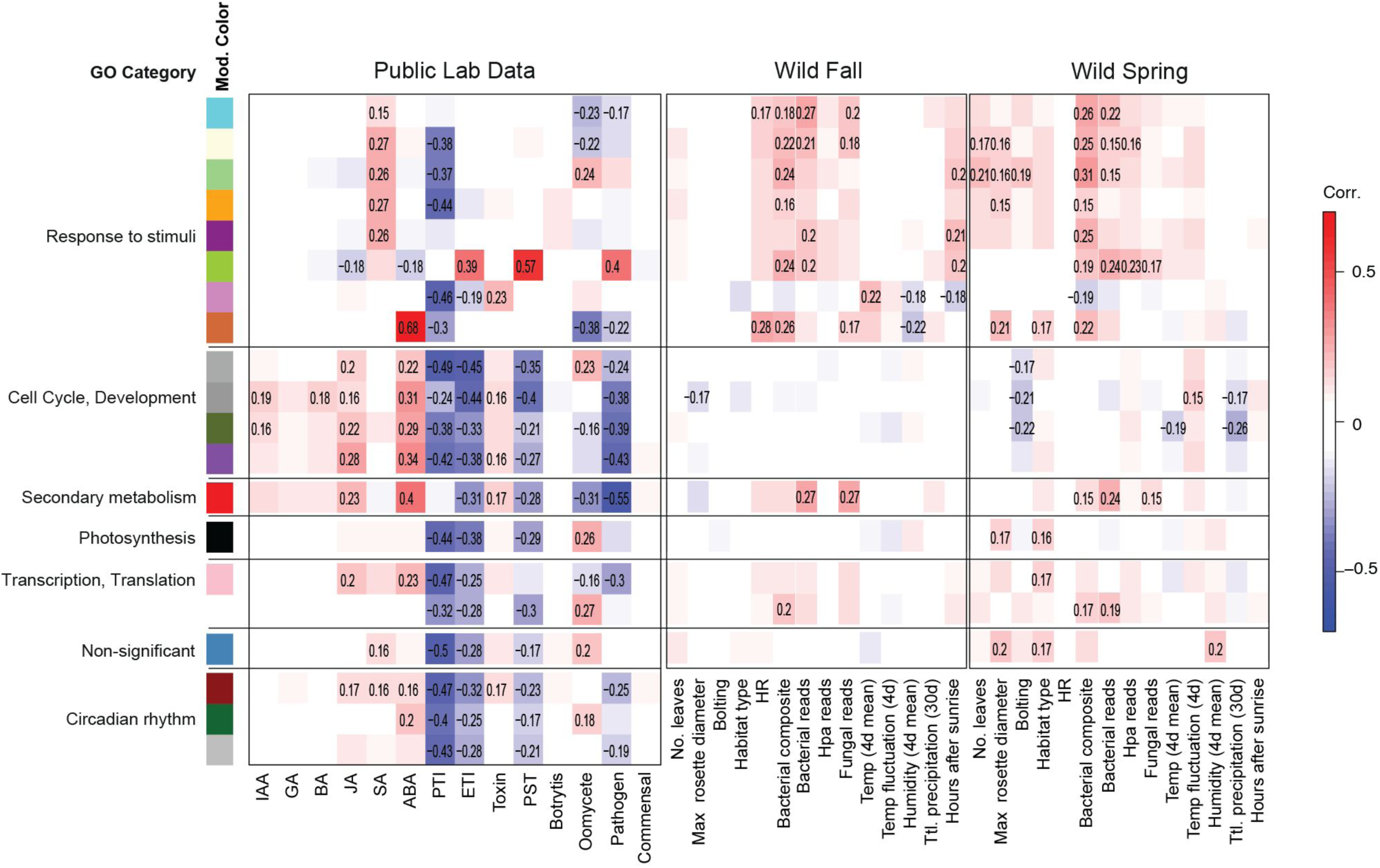
Module–trait correlation heatmaps. Pearson correlations between eigengenes (MEs) of the public biotic WGCNA and laboratory treatments (left), and between eigengenes of wild fall (middle) and spring (right) transcriptomes projected onto the biotic modules and their corresponding field scores. Module colors are indicated by rectangles on the left, and modules are grouped into six GO categories. Numbers in the heatmaps show Pearson’s *r* for significant ME–trait associations only (*p* < 0.005). Lab MEs exhibit strong, bidirectional correlations with applied treatments, whereas associations between projected wild MEs and field scores are weaker but generally positive. The strong negative associations between laboratory infections and growth-related functions are most notably missing in the wild.

**Extended Figure 3.1:**
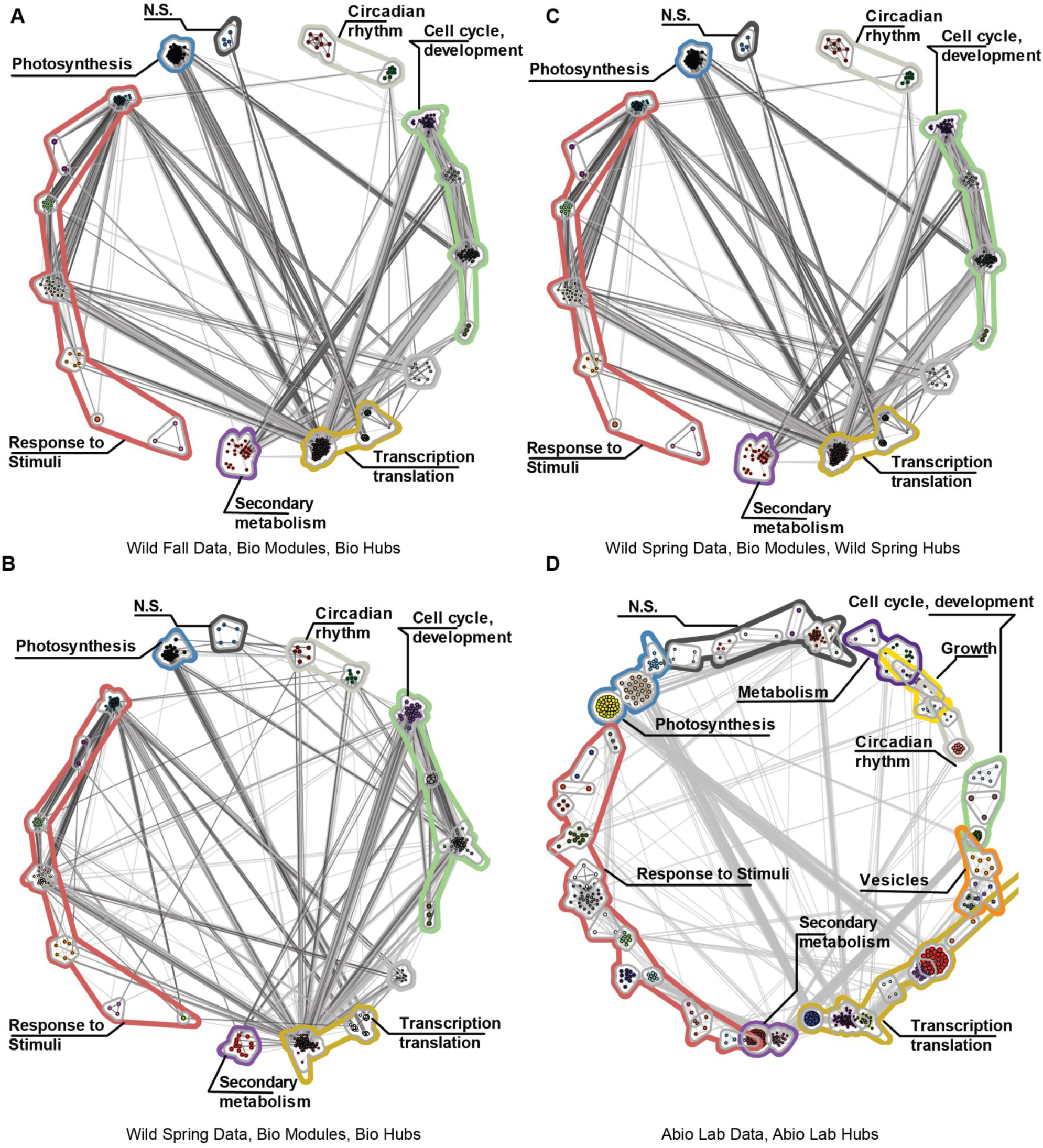
Hub-gene networks from wild-projected biotic networks (A-C) and abiotic lab networks (D). **A-B.** Hub-gene networks constructed from hub genes (top 10% by within-module connectivity) identified in biotic networks projected from wild samples collected in fall 2022 (A) and spring 2023 (B). **C.** Network constructed from hub genes identified within the lab biotic network, with inter-gene relationships projected from

**Extended Figure 3.2:**
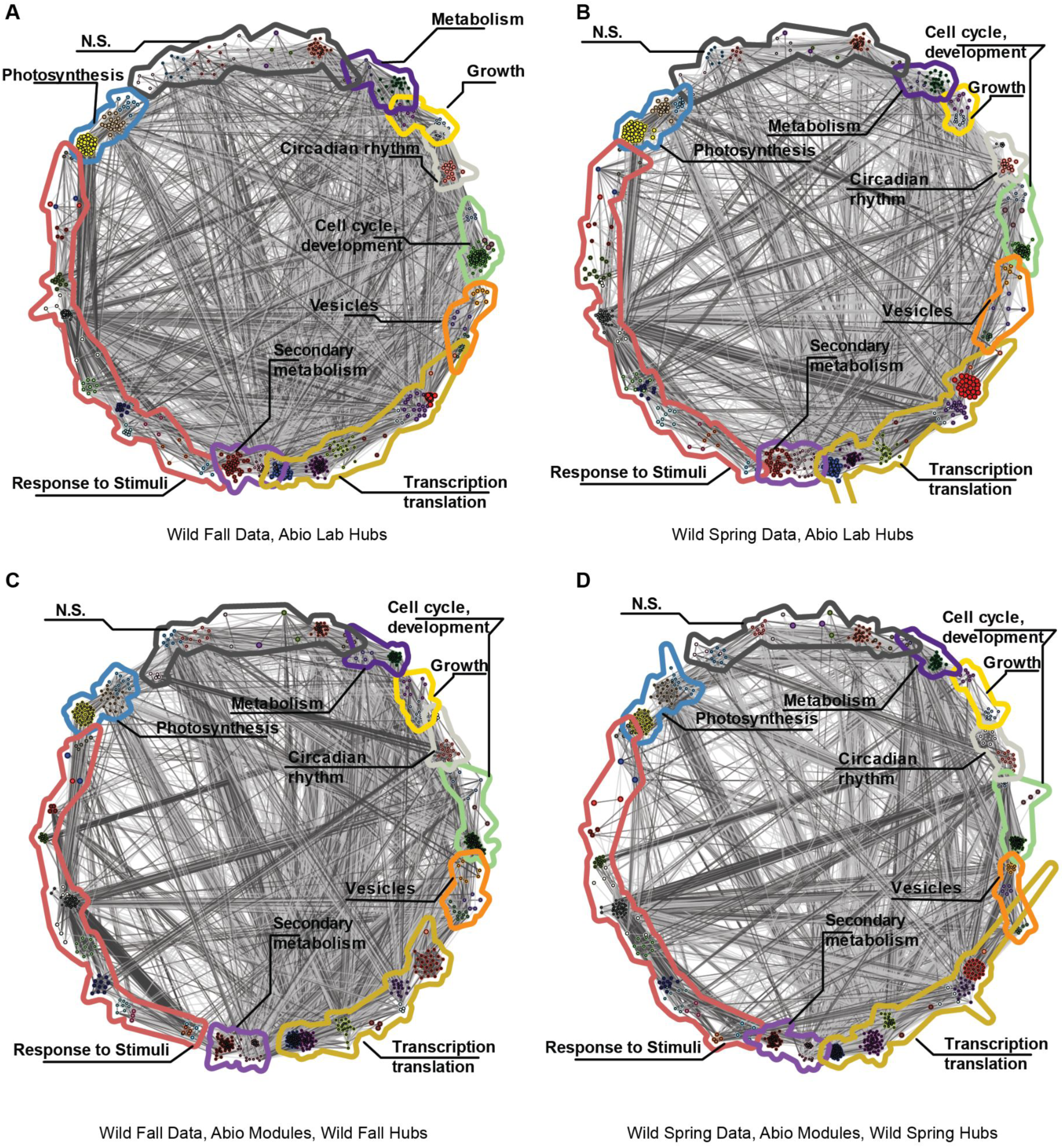
Hub-gene networks from wild-projected abiotic networks. **A-B.** Hub-gene networks constructed from hub genes identified within the lab abiotic network, with inter-gene relationships projected from

## Supplementary information

### Supplementary Figures

**Figure. S1:**
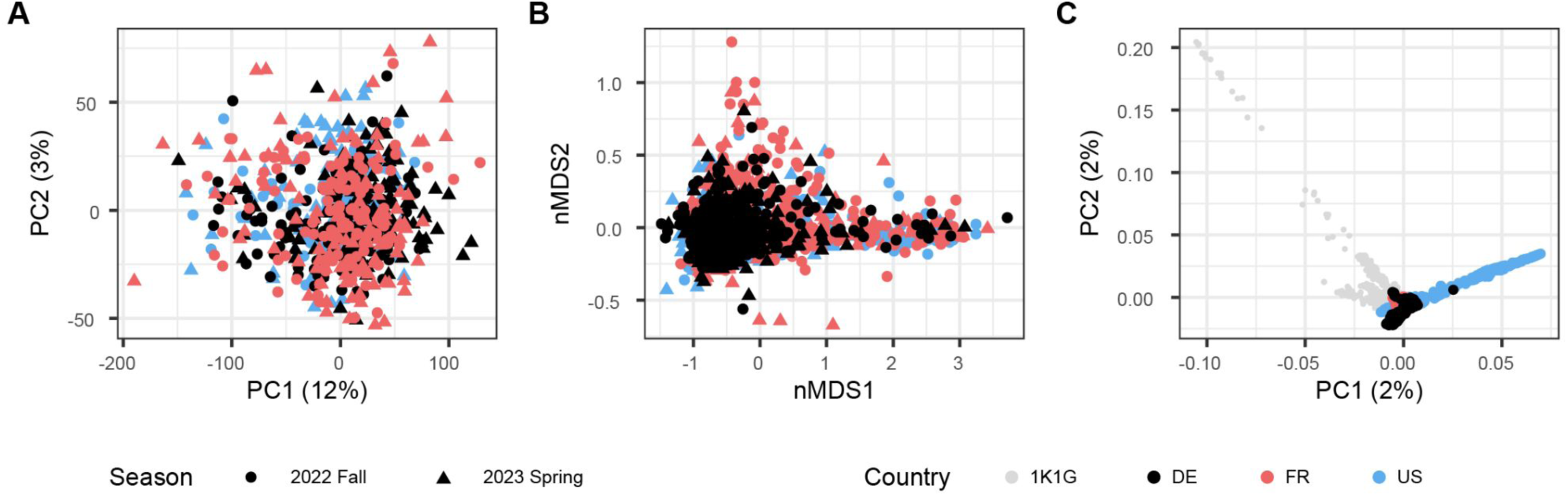
Distribution of A) transcriptomic, B) metagenomic, and C) host genetic variation among samples. While host genetics is structured by sampling location, transcriptomes and microbiomes are not structured by either country or season.

**Figure S2:**
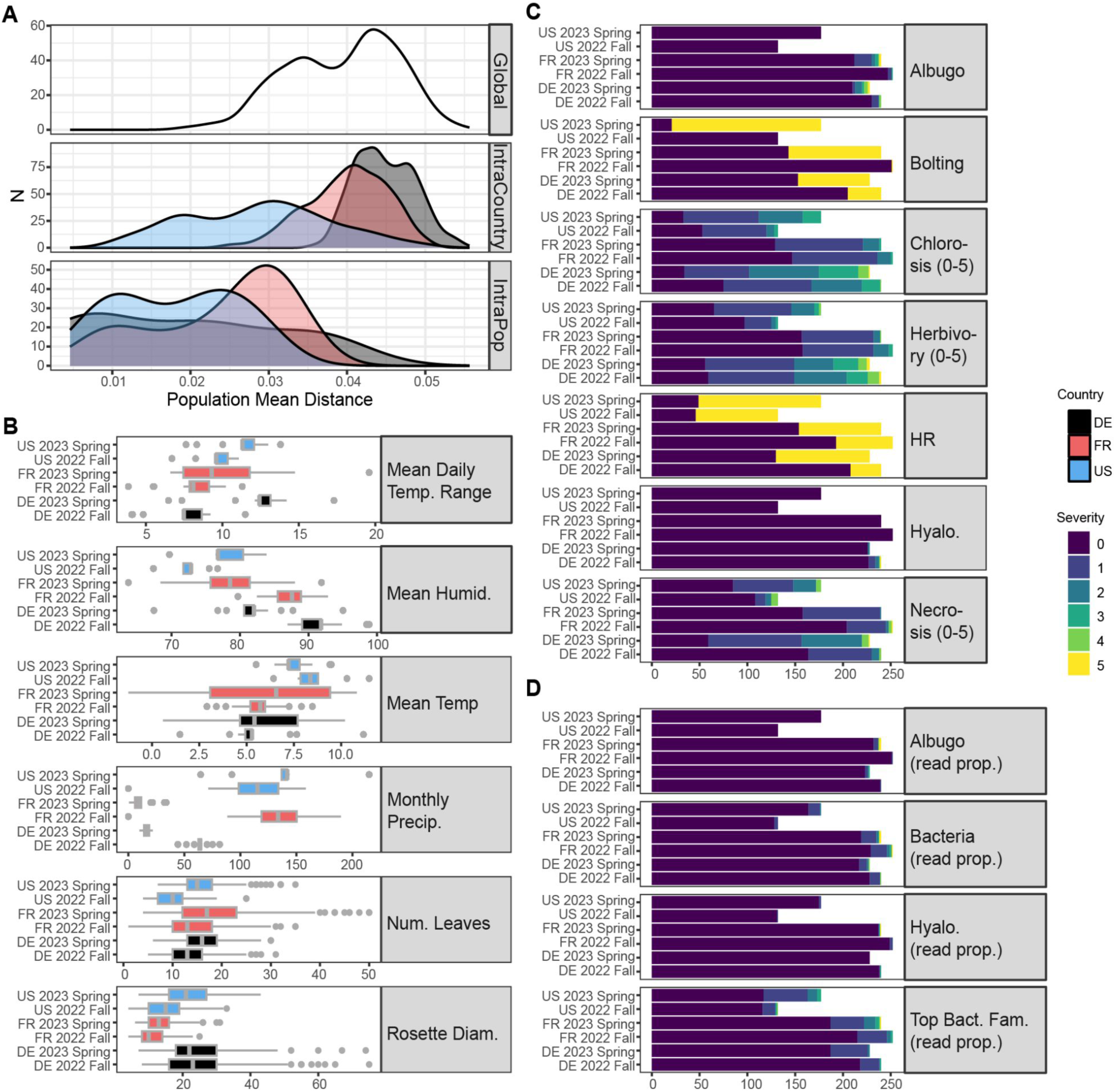
Distribution of genetic, developmental, and disease characters across all plants. A) Histograms of inter-sample distances for plant pairs within sites (IntraPop), within countries but between sites (IntraCountry) and between countries (Global). For DE/FR, intra site distances are considerably smaller than inter-site, while USA has generally lower diversity. B) Distribution of weather and plant growth among countries and seasons. While we made efforts to ensure collections occurred at comparable climatic conditions and developmental stages between countries, some differences between countries remain, due to seasonal variation in weather over the growing

**Figure S3:**
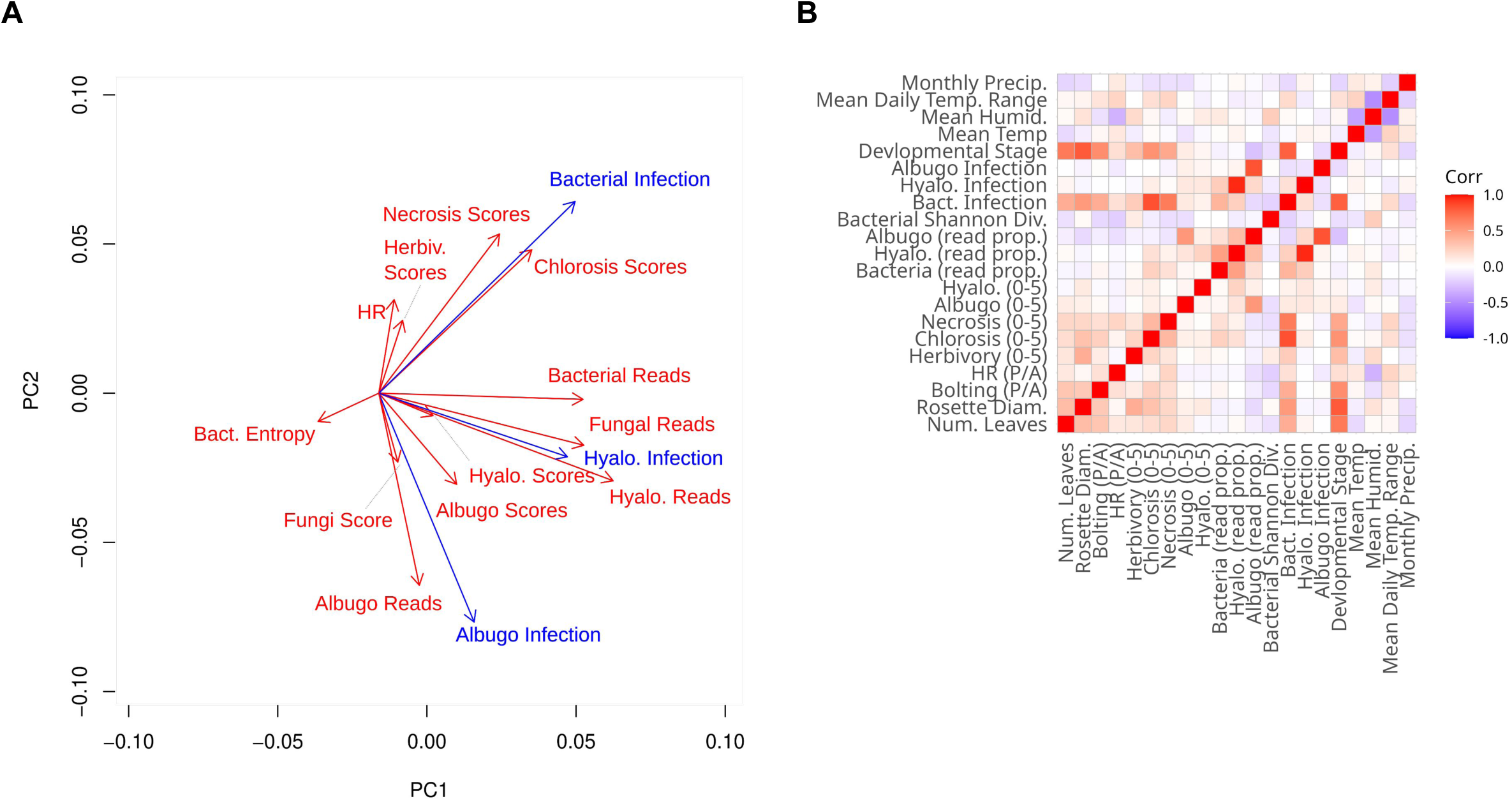
Confirmatory factor analysis to synthesize latent variable representations of bacterial, hyaloperonospora, and albugo infections from multiple imprecise observations. **A)** PCA highlights correlations among partially redundant measures of microbial infection. Latent estimates of infection (blue) integrated across these multiple predictors to form simple estimates of infection. **B)** Correlation among weather, microbial load, and plant disease scores and size. Albugo scores and load are highly positively correlated, while Hyaloperonospora and bacterial loads have weaker positive correlations.

**Figure S4:**
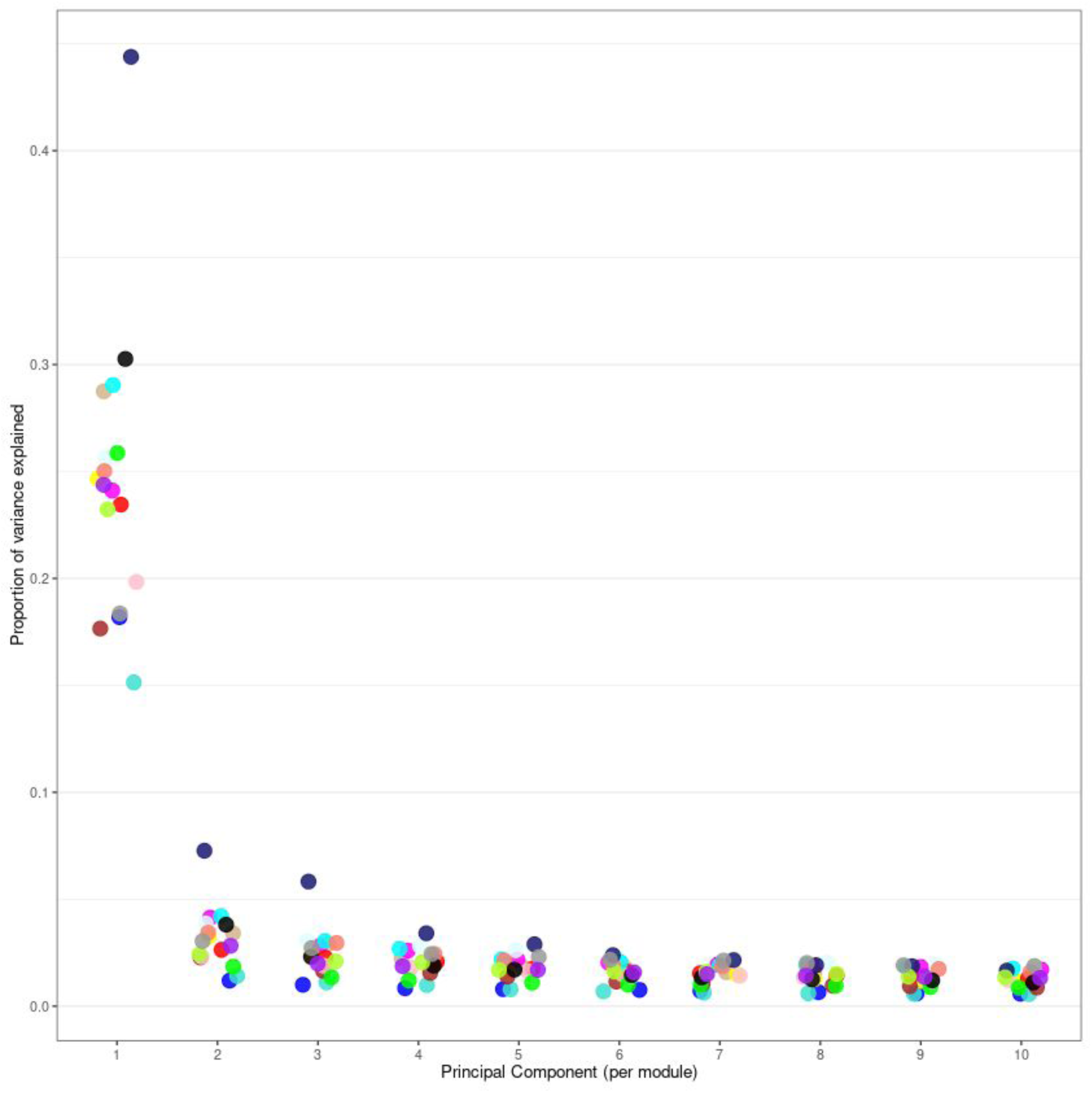
Proportion of variance explained by the first 10 module PCs of the wild RNA data. Each point represents one module, coloured by their respective module identity. PC1 corresponds to the module eigengene (ME), which explained between 15-45% of the variance in respective modules.

**Figure S5:**
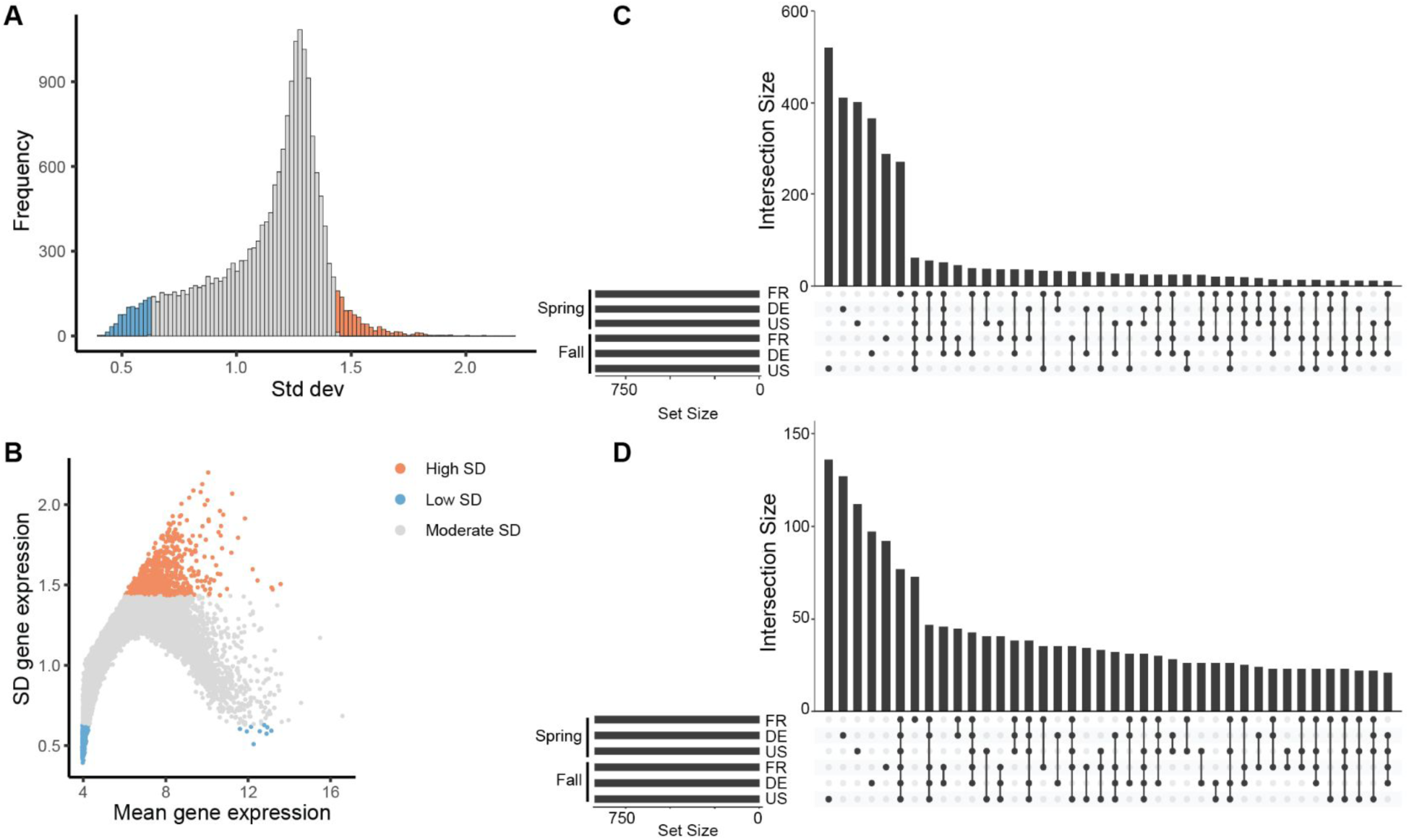
Characterization of the high- and low-SD genes. **A.** Histogram of standard deviation of all genes passing initial dropout and variance filter. Genes ranking top and bottom 5% of SD was identified as high- and low- SD genes (N=922 each). **B.** Relationships between average gene expression level and its SD. Note that while low SD genes occupy expression extremes, high SD genes all have moderate expression levels. **C-D**. Upset plot showing gene sharing among high (panel C) and low (panel D) SD genes. High- and low-SD genes were called separately for each study region and season. The great majority of high- and low-SD genes are private to each region and season, especially for high SD genes.

**Figure S6:**
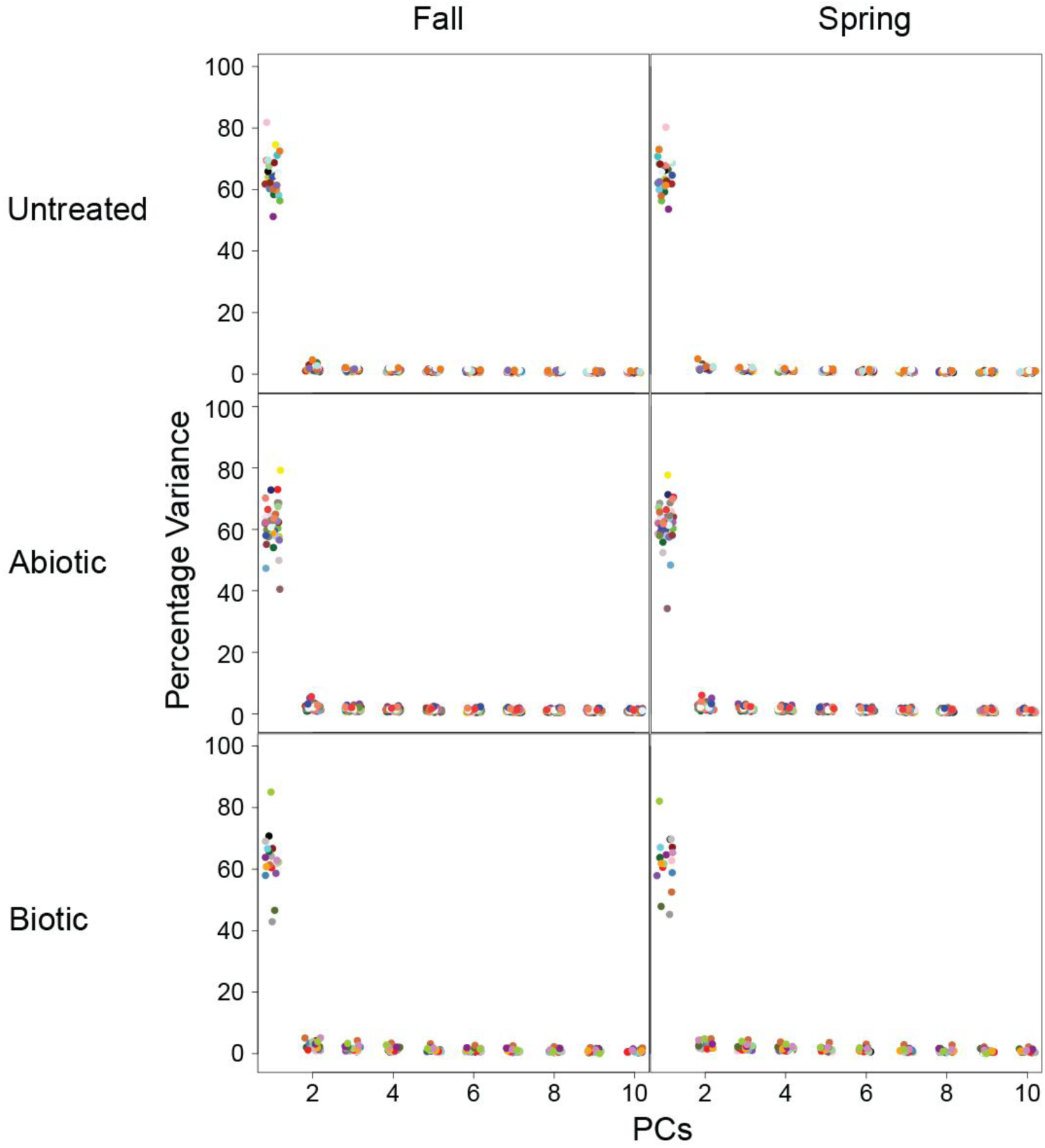
Proportion of variance explained by the first 10 module PCs of the wild-projected lab WGCNs. Module assignments were acquired from independent WGCNA using published lab data of untreated (top), abiotic (middle), and biotic (bottom) treatment samples, and wild expression data from fall 2022 (left) and spring 2023 (right) were projected onto the modules. Each point represents one module, colored according to their designated color labels. PC1 corresponds to the module eigengene (ME), which explained an average of ∼60% of the variance in respective modules.

**Figure S7.**
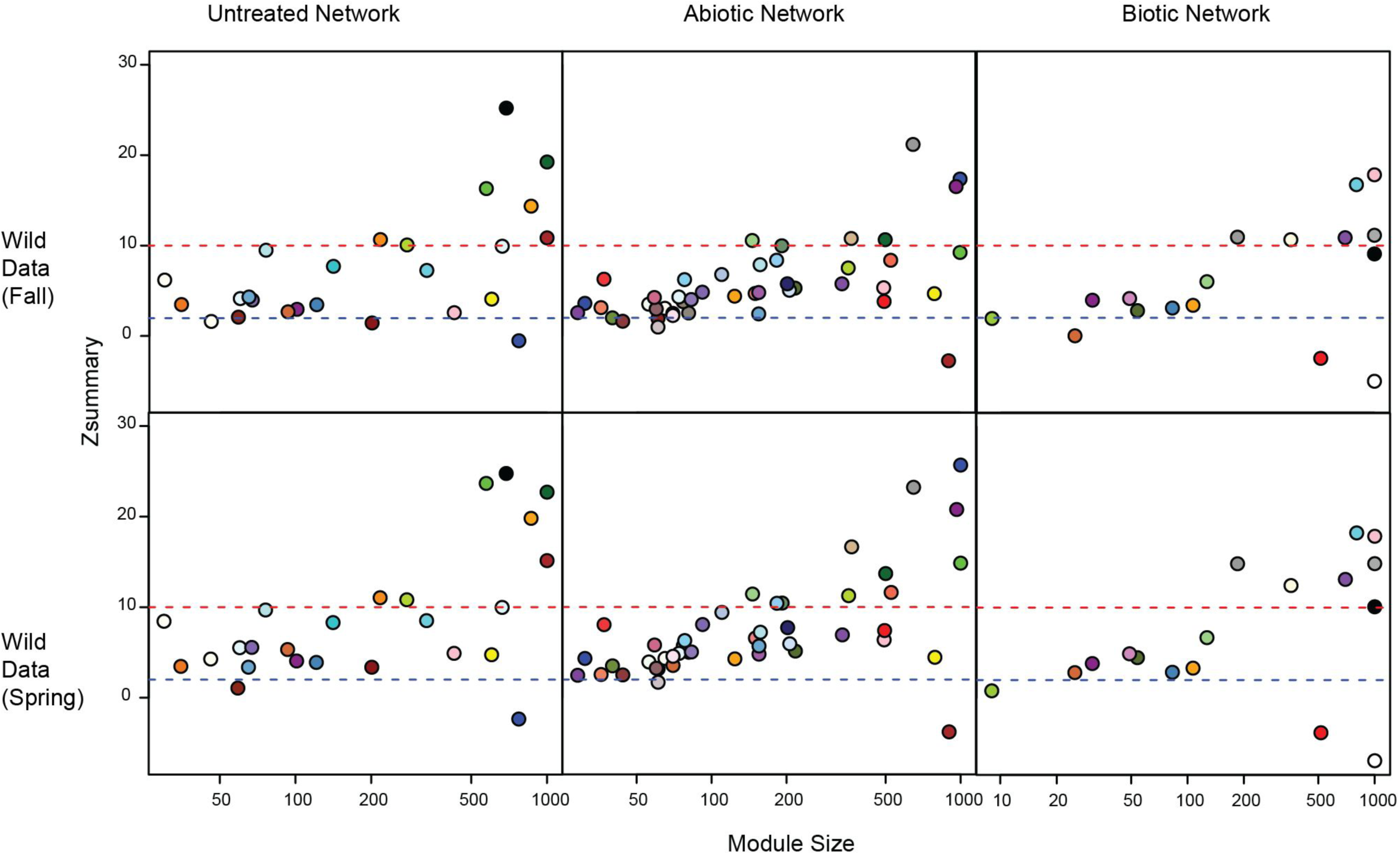
Module preservation statistics of lab-derived WGCNs after the projection of wild fall (top panel) and spring (bottom panel) samples. Modules are colored according to designated color labels. Z-summary score>10: highly conserved, >2: moderately conserved, <2: unconserved. Both untreated and abiotic projections showed abundant moderately conserved modules, while biotic network has less middle category, and featured greater proportion of highly conserved modules.

**Figure S8:**
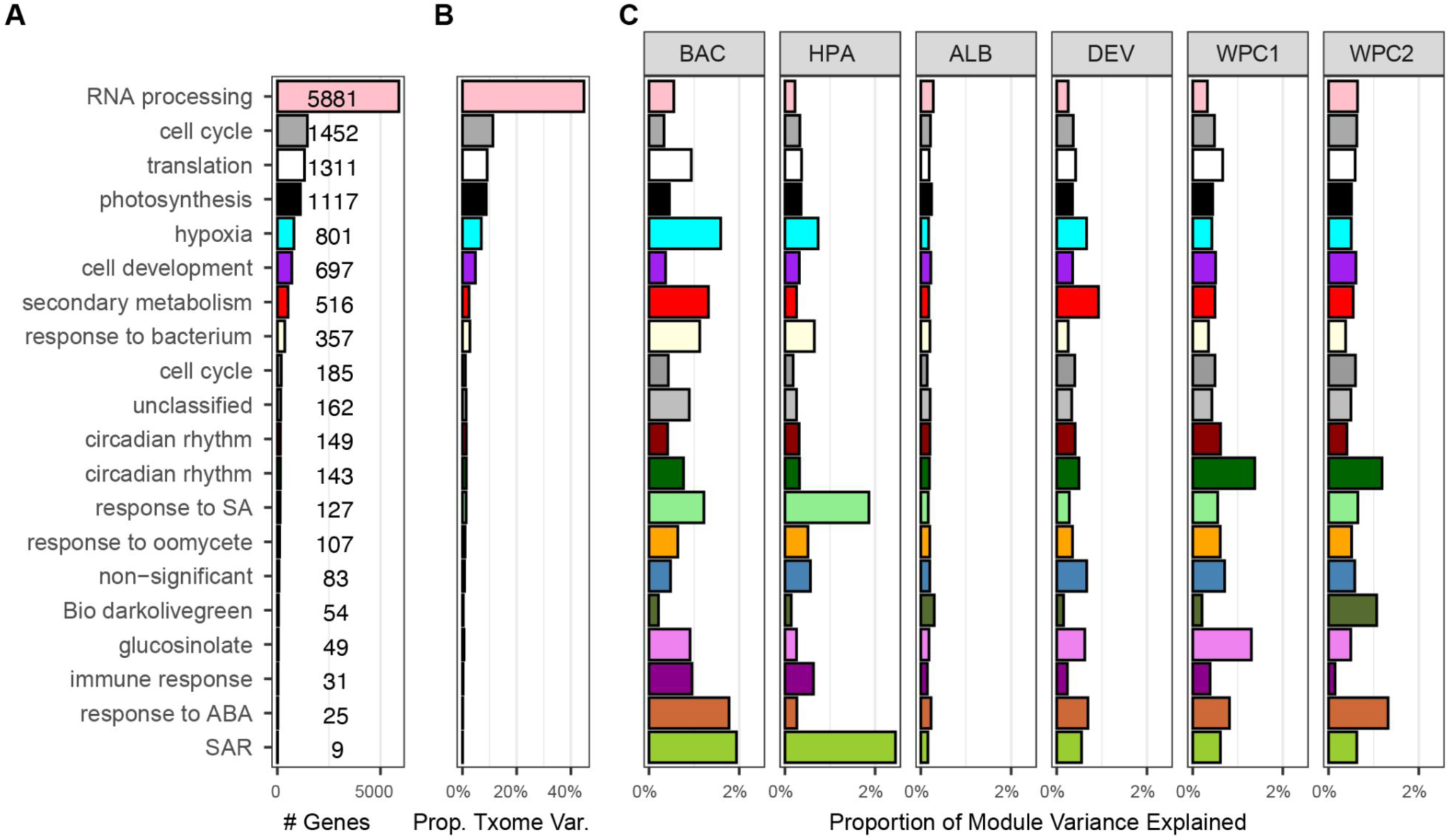
RDA variance decomposition across biotic lab modules. A) Proportion of global transcriptome variance attributable to each of the projected biotic lab WCGNA modules, and number of genes in each module. B) Proportion of module expression variance explained by constraints in per-module RDA analyses, constrained by the same factors as main Fig 3. C) Proportion global module expression variance uniquely explained by each predictor.

**Figure S9:**
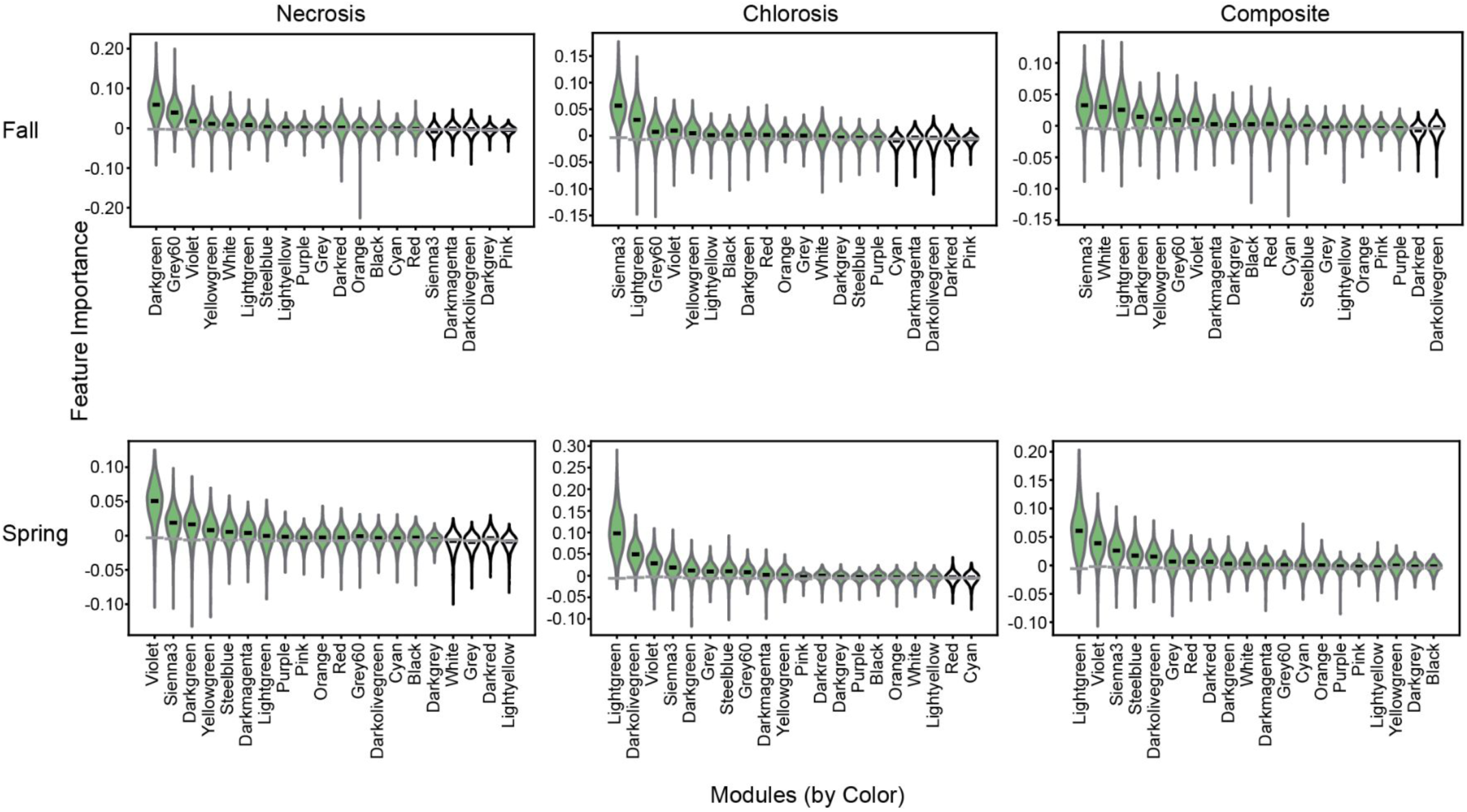
Feature importance scores for each module eigengene (ME) in random forest predictions. Necrosis (left), chlorosis (middle), and their composite (right) from fall (top) and spring (bottom) data, respectively. MEs were calculated after projection of wild data onto public biotic modules. Each violin shows the distribution of module feature importance in 1000x random forest iterations, while the grey bars at the bottom of each violin illustrates the median of 1000x permuted null distribution.

## References

Arana, María Verónica, and F. Xavier Picó. 2025. “Advancing Ecological and Evolutionary Research in *Arabidopsis*: Extending Insights into Model and Nonmodel Plants.” The Plant Cell 37 (7): koaf151.

Atwell, Susanna, Yu S. Huang, Bjarni J. Vilhjálmsson, et al. 2010. “Genome-Wide Association Study of 107 Phenotypes in *Arabidopsis thaliana* Inbred Lines.” Nature 465 (7298): 627–631.

Bai, Yang, Daniel B. Müller, Girish Srinivas, et al. 2015. “Functional Overlap of the *Arabidopsis* Leaf and Root Microbiota.” Nature 528 (7582): 364–369.

Bartoli, Claudia, Léa Frachon, Matthieu Barret, et al. 2018. “In Situ Relationships between Microbiota and Potential Pathobiota in *Arabidopsis* thaliana.” The ISME Journal 12 (8): 2024–2038.

Baskin, Jerry M., and Carol C. Baskin. 1983. “Seasonal Changes in the Germination Responses of Buried Seeds of *Arabidopsis* thaliana and Ecological Interpretation.” *Botanical Gazette (Chicago*, Ill.) 144 (4): 540–543.

Bergelson, J., and C. B. Purrington. 1996. “Surveying Patterns in the Cost of Resistance in Plants.” The American Naturalist 148 (3): 536–558.

Bhaskara, Govinal Badiger, Taslima Haque, Jason E. Bonnette, et al. 2023. “Evolutionary Analyses of Gene Expression Divergence in Panicum Hallii: Exploring Constitutive and Plastic Responses Using Reciprocal Transplants.” Molecular Biology and Evolution 40 (10). 10.1093/molbev/msad210.

Bojanowski, Michał. 2023. “Intergraph: Coercion Routines for Network Data Objects.” https://mbojan.github.io/intergraph/.

Bomblies, Kirsten, Levi Yant, Roosa A. Laitinen, et al. 2010. “Local-Scale Patterns of Genetic Variability, Outcrossing, and Spatial Structure in Natural Stands of *Arabidopsis* thaliana.” PLoS Genetics 6 (3): e1000890.

Brachi, B., N. Faure, M. Horton, et al. 2010. “Linkage and Association Mapping of *Arabidopsis* thaliana Flowering Time in Nature.” PLoS Genetics 6: e1000940.

Brown, J. K. 2002. “Yield Penalties of Disease Resistance in Crops.” Current Opinion in Plant Biology 5 (4): 339–344.

Butts, Carter T. 2008. “Network: A Package for Managing Relational Data in R.” Journal of Statistical Software 24 (2): 1–36.

Carlson, Marc. 2024. “org.At.tair.db: Genome Wide Annotation for *Arabidopsis*.” Preprint.

Cheng, Chia-Yi, Vivek Krishnakumar, Agnes P. Chan, Françoise Thibaud-Nissen, Seth Schobel, and Christopher D. Town. 2017. “Araport11: A Complete Reannotation of the *Arabidopsis* thaliana Reference Genome.” The Plant Journal: For Cell and Molecular Biology 89 (4): 789–804.

Conesa, Ana, Pedro Madrigal, Sonia Tarazona, et al. 2016. “A Survey of Best Practices for RNA-Seq Data Analysis.” Genome Biology 17 (January): 13.

Cook, Di, Joseph Larmarange, Francois Briatte, et al. 2025. “GGally: Extension to ‘ggplot2.’” Comprehensive R Archive Network (CRAN), August 23. https://CRAN.R-project.org/package=GGally.

Csárdi, Gábor, Tamás Nepusz, Vincent Traag, et al. 2026. “Igraph: Network Analysis and Visualization in R.” Preprint. 10.5281/zenodo.7682609.

Des Marais, David L., Kyle M. Hernandez, and Thomas E. Juenger. 2013. “Genotype-by-Environment Interaction and Plasticity: Exploring Genomic Responses of Plants to the Abiotic Environment.” Annual Review of Ecology, Evolution, and Systematics 44 (1): 5–29.

Dixon, Philip. 2003. “VEGAN, a Package of R Functions for Community Ecology.” Journal of Vegetation Science: Official Organ of the International Association for Vegetation Science 14 (6): 927–930.

Donohue, Kathleen, Rafael Rubio de Casas, Liana Burghardt, Katherine Kovach, and Charles G. Willis. 2010. “Germination, Postgermination Adaptation, and Species Ecological Ranges.” Annual Review of Ecology, Evolution, and Systematics 41 (1): 293–319.

Duque-Jaramillo, Alejandra, Nina Ulmer, Saleh Alseekh, et al. 2023. “The Genetic and Physiological Basis of *Arabidopsis* thaliana Tolerance to Pseudomonas Viridiflava.” *The New Phytologist*, ahead of print, September 4. 10.1111/nph.19241.

Fournier-Level, A., A. Korte, Cooper, M. Nordborg, J. Schmitt, and A. M. Wilczek. 2011. “A Map of Local Adaptation in *Arabidopsis* thaliana.” Science, 1–27.

Frachon, Léa, Claudia Bartoli, Sébastien Carrère, et al. 2018. “A Genomic Map of Climate Adaptation in *Arabidopsis* thaliana at a Micro-Geographic Scale.” Frontiers in Plant Science 9. 10.3389/fpls.2018.00967.

Goss, Erica M., and Joy Bergelson. 2007. “Fitness Consequences of Infection of *Arabidopsis* thaliana with Its Natural Bacterial Pathogen Pseudomonas Viridiflava.” Oecologia 152 (1): 71–81.

Groen, Simon C., Irina Ćalić, Zoé Joly-Lopez, et al. 2020. “The Strength and Pattern of Natural Selection on Gene Expression in Rice.” Nature 578 (7796): 572–576.

Gurung, Priya Darshini, Atul Kumar Upadhyay, Pardeep Kumar Bhardwaj, Ramanathan Sowdhamini, and Uma Ramakrishnan. 2019. “Transcriptome Analysis Reveals Plasticity in Gene Regulation due to Environmental Cues in Primula Sikkimensis, a High Altitude Plant Species.” BMC Genomics 20 (1): 989.

Jakob, Katrin, Erica M. Goss, Hitoshi Araki, Tam Van, Martin Kreitman, and Joy Bergelson. 2002. “Pseudomonas Viridiflava and P. Syringae--Natural Pathogens of *Arabidopsis* thaliana.” Molecular Plant-Microbe Interactions: MPMI 15 (12): 1195–1203.

Korves, Tonia, and Joy Bergelson. 2004. “A Novel Cost of R Gene Resistance in the Presence of Disease.” The American Naturalist 163 (4): 489–504.

Köster, Johannes, and Sven Rahmann. 2012. “Snakemake --- a Scalable Bioinformatics Workflow Engine.” Bioinformatics 28 (19): 2520–2522.

Lamesch, Philippe, Tanya Z. Berardini, Donghui Li, et al. 2012. “The *Arabidopsis* Information Resource (TAIR): Improved Gene Annotation and New Tools.” Nucleic Acids Research 40 (Database issue): D1202–10.

Lämke, Jörn, and Isabel Bäurle. 2017. “Epigenetic and Chromatin-Based Mechanisms in Environmental Stress Adaptation and Stress Memory in Plants.” Genome Biology 18 (1): 124.

Langfelder, Peter, and Steve Horvath. 2008. “WGCNA: An R Package for Weighted Correlation Network Analysis.” BMC Bioinformatics 9 (1): 559.

Lempe, Janne, Sureshkumar Balasubramanian, Sridevi Sureshkumar, Anandita Singh, Markus Schmid, and Detlef Weigel. 2005. “Diversity of Flowering Responses in Wild *Arabidopsis* thaliana Strains.” PLoS Genetics 1 (1): 109–118.

Li, Bo, and Colin N. Dewey. 2011. “RSEM: Accurate Transcript Quantification from RNA-Seq Data with or without a Reference Genome.” BMC Bioinformatics 12 (1): 323.

Li, Heng. 2008. “Seqtk - Toolkit for Processing Sequences in FASTA/Q Formats.” In GitHub. Preprint. https://github.com/lh3/seqtk.

Li, Heng. 2013. Aligning Sequence Reads, Clone Sequences and Assembly Contigs with BWA-MEM. March.

Loewa, Anna, James J. Feng, and Sarah Hedtrich. 2023. “Human Disease Models in Drug Development.” Nature Reviews Bioengineering 1 (8): 1–15.

Love, Michael I., Wolfgang Huber, and Simon Anders. 2014. “Moderated Estimation of Fold Change and Dispersion for RNA-Seq Data with DESeq2.” Genome Biology 15 (12): 550.

Lundberg, Derek S., Joy Bergelson, Fabrice Roux, Detlef Weigel, and Talia L. Karasov. 2025. “Lab to Field: Challenges and Opportunities for Plant Biology.” Cell Host & Microbe 33 (8): 1212–1216.

Lundberg, Derek S., Sonja Kersten, Ezgi Mehmetoğlu Boz, et al. 2025. “A Major Trade-off between Growth and Defense in *Arabidopsis* thaliana Can Vanish in Field Conditions.” PLoS Biology 23 (7): e3003237.

MacQueen, Alice, Xiaoqin Sun, and Joy Bergelson. 2016. “Genetic Architecture and Pleiotropy Shape Costs of Rps2-Mediated Resistance in *Arabidopsis* thaliana.” Nature Plants 2 (July): 16110.

Maier, Benjamin A., Patrick Kiefer, Christopher M. Field, et al. 2021. “A General Non-Self Response as Part of Plant Immunity.” Nature Plants 7 (5): 696–705.

Mauricio, R. 1998. “Costs of Resistance to Natural Enemies in Field Populations of the Annual Plant *Arabidopsis* thaliana.” The American Naturalist 151 (1): 20–28.

Mi, Huaiyu, Anushya Muruganujan, Dustin Ebert, Xiaosong Huang, and Paul D. Thomas. 2019. “PANTHER Version 14: More Genomes, a New PANTHER GO-Slim and Improvements in Enrichment Analysis Tools.” Nucleic Acids Research 47 (D1): D419–D426.

Mjema, Eneza Yoeli, Maria Letícia Bonatelli, Dirk C. Albach, et al. 2026. “Molecular and Phenotypic Footprints of Climate in Native Arabidopsis *thaliana*.” In *bioRxiv*. BioRxiv, March 4. 10.64898/2026.03.02.709013.

Murray, Kevin D., Justin O. Borevitz, Detlef Weigel, and Norman Warthmann. 2024. “Acanthophis: A Comprehensive Plant Hologenomics Pipeline.” Journal of Open Source Software 9 (95): 6062.

Nagano, Atsushi J., Yutaka Sato, Motohiro Mihara, et al. 2012. “Deciphering and Prediction of Transcriptome Dynamics under Fluctuating Field Conditions.” Cell 151 (6): 1358–1369.

Nagano, Atsushi J., Tetsuhiro Kawagoe, Jiro Sugisaka, Mie N. Honjo, Koji Iwayama, and Hiroshi Kudoh. 2019. “Annual Transcriptome Dynamics in Natural Environments Reveals Plant Seasonal Adaptation.” Nature Plants 5 (1): 74–83.

Ojha, Megha, Dilip G. T. Naidu, and Sumanta Bagchi. 2022. “Meta-analysis of Induced Anti-herbivore Defence Traits in Plants from 647 Manipulative Experiments with Natural and Simulated Herbivory.” The Journal of Ecology 110 (4): 799–816.

Pedregosa, Fabian, Gaël Varoquaux, Alexandre Gramfort, et al. 2011. “Scikit-Learn: Machine Learning in Python.” Journal of Machine Learning Research 12 (85): 2825–2830.

Picelli, Simone, Asa K. Björklund, Björn Reinius, Sven Sagasser, Gösta Winberg, and Rickard Sandberg. 2014. “Tn5 Transposase and Tagmentation Procedures for Massively Scaled Sequencing Projects.” Genome Research 24 (12): 2033–2040.

Plessis, Anne, Christoph Hafemeister, Olivia Wilkins, et al. 2015. “Multiple Abiotic Stimuli Are Integrated in the Regulation of Rice Gene Expression under Field Conditions.” eLife 4 (November). 10.7554/eLife.08411.

Poplin, Ryan, Pi-Chuan Chang, David Alexander, et al. 2018. “A Universal SNP and Small-Indel Variant Caller Using Deep Neural Networks.” Nature Biotechnology 36 (10): 983–987.

R Core Team. 2024. “R: A Language and Environment for Statistical Computing.” Preprint, R Foundation for Statistical Computing. https://www.R-project.org/.

Regalado, Julian, Derek S. Lundberg, Oliver Deusch, et al. 2020. “Combining Whole-Genome Shotgun Sequencing and rRNA Gene Amplicon Analyses to Improve Detection of Microbe\textendash Microbe Interaction Networks in Plant Leaves.” The ISME Journal 14 (8): 2116–2130.

Richards, Christina L., Ulises Rosas, Joshua Banta, Naeha Bhambhra, and Michael D. Purugganan. 2012. “Genome-Wide Patterns of *Arabidopsis* Gene Expression in Nature.” PLoS Genetics 8 (4): e1002662.

Ritchie, Matthew E., Belinda Phipson, Di Wu, et al. 2015. “Limma Powers Differential Expression Analyses for RNA-Sequencing and Microarray Studies.” Nucleic Acids Research 43 (7): e47.

Rosseel, Yves. 2012. “Lavaan: An R Package for Structural Equation Modeling.” Journal of Statistical Software.

Roux, Fabrice, and Léa Frachon. 2022. “A Genome-Wide Association Study in *Arabidopsis* thaliana to Decipher the Adaptive Genetics of Quantitative Disease Resistance in a Native Heterogeneous Environment.” PloS One 17 (10): e0274561.

Schubert, Mikkel, Stinus Lindgreen, and Ludovic Orlando. 2016. “AdapterRemoval v2: Rapid Adapter Trimming, Identification, and Read Merging.” BMC Research Notes 9: 88.

Shirsekar, Gautam, Jane Devos, Sergio M. Latorre, et al. 2021. “Multiple Sources of Introduction of North American *Arabidopsis* thaliana from across Eurasia.” Molecular Biology and Evolution 38 (12): 5328–5344.

Smyth, G. K. 2005. “Limma: Linear Models for Microarray Data.” In Bioinformatics and Computational Biology Solutions Using R and Bioconductor. Springer-Verlag.

Stergiopoulos, Ioannis, and Thomas R. Gordon. 2014. “Cryptic Fungal Infections: The Hidden Agenda of Plant Pathogens.” Frontiers in Plant Science 5 (September): 506.

Taguas, Ignacio, François Maclot, Nuria Montes, Israel Pagán, Aurora Fraile, and Fernando García-Arenal. 2025. “Infection Patterns of Albugo Laibachii and Effect on Host Survival and Reproduction in a Wild Population of *Arabidopsis* thaliana.” Plants 14 (4): 568.

Teasdale, Luisa C., Kevin D. Murray, Max Collenberg, et al. 2025. “Pangenomic Context Reveals the Extent of Intraspecific Plant NLR Evolution.” Cell Host & Microbe 33 (8): 1291–1305.e9.

Thines, M., Y-J Choi, E. Kemen, et al. 2009. “A New Species of Albugo Parasitic to *Arabidopsis* thaliana Reveals New Evolutionary Patterns in White Blister Rusts (Albuginaceae).” Persoonia 22 (1): 123–128.

Todesco, Marco, Sureshkumar Balasubramanian, Tina T. Hu, et al. 2010. “Natural Allelic Variation Underlying a Major Fitness Trade-off in *Arabidopsis* thaliana.” Nature 465 (7298): 632–636.

Walter, Greg M., James Clark, Delia Terranova, et al. 2023. “Hidden Genetic Variation in Plasticity Provides the Potential for Rapid Adaptation to Novel Environments.” The New Phytologist 239 (1): 374–387.

Weinig, C., L. A. Dorn, N. C. Kane, et al. 2003. “Heterogeneous Selection at Specific Loci in Natural Environments in *Arabidopsis* thaliana.” Genetics 165 (1): 321–329.

Weinig, C., J. R. Stinchcombe, and J. Schmitt. 2003. “QTL Architecture of Resistance and Tolerance Traits in *Arabidopsis* thaliana in Natural Environments.” Molecular Ecology 12 (5): 1153–1163.

Wersch, Rowan van, Xin Li, and Yuelin Zhang. 2016. “Mighty Dwarfs: *Arabidopsis* Autoimmune Mutants and Their Usages in Genetic Dissection of Plant Immunity.” Frontiers in Plant Science 7 (November): 1717.

Wickham, Hadley. 2016. Ggplot2. 2nd ed. Use R! Springer International Publishing. PDF.

Wlodzimierz, Piotr, Fernando A. Rabanal, Robin Burns, et al. 2023. “Cycles of Satellite and Transposon Evolution in *Arabidopsis* Centromeres.” Nature 618 (7965): 557–565.

Wood, Derrick E., Jennifer Lu, and Ben Langmead. 2019. “Improved Metagenomic Analysis with Kraken 2.” Genome Biology 20 (1): 257.

Yaffe, Hila, Kobi Buxdorf, Illil Shapira, et al. 2012. “LogSpin: A Simple, Economical and Fast Method for RNA Isolation from Infected or Healthy Plants and Other Eukaryotic Tissues.” BMC Research Notes 5 (January): 45.

Yuan, Wei, Fiona Beitel, Thanvi Srikant, et al. 2023. “Pervasive under-Dominance in Gene Expression Underlying Emergent Growth Trajectories in *Arabidopsis* thaliana Hybrids.” Genome Biology 24 (1): 200.

Yu, Guangchuang. 2024. “Thirteen Years of clusterProfiler.” Innovation (Cambridge (Mass.)) 5 (6): 100722.

Yun, Taedong, Helen Li, Pi-Chuan Chang, Michael F. Lin, Andrew Carroll, and Cory Y. McLean. 2021. “Accurate, Scalable Cohort Variant Calls Using DeepVariant and GLnexus.” *Bioinformatics (Oxford*, England) 36 (24): 5582–5589.

Yu, Yiming, Hong Zhang, Yanping Long, Yi Shu, and Jixian Zhai. 2022. “Plant Public RNA-Seq Database: A Comprehensive Online Database for Expression Analysis of ∼45 000 Plant Public RNA-Seq Libraries.” Plant Biotechnology Journal 20 (5): 806–808.

Züst, Tobias, and Anurag A. Agrawal. 2017. “Trade-Offs between Plant Growth and Defense against Insect Herbivory: An Emerging Mechanistic Synthesis.” Annual Review of Plant Biology 68 (1): 513–534.

